# Personalized microbiotas (counter-)select for antibiotic resistant pathogens

**DOI:** 10.64898/2026.03.29.715108

**Authors:** Michael Knopp, Sarela Garcia-Santamarina, Lina Michel, Dimitrios Papagiannidis, Sophia David, Denise M. Selegato, Joshua L. C. Wong, Nicolai Karcher, Gad Frankel, Michael Zimmermann, Mikhail M. Savitski, Athanasios Typas

## Abstract

Antibiotic resistant pathogens are an increasing public health threat, as development of novel therapeutics is outpaced by resistance emergence and dissemination. Approaches to slow down or even revert antibiotic resistance are necessary to maintain efficacy of both existing and new antibiotics. Such approaches exploit the fitness cost of resistance elements, but have largely relied on assessing this cost in laboratory conditions that poorly reflect the native context in which pathogens reside. Here we present a method that allows to investigate the influence of personalized human gut microbiota compositions on the competitive fitness of antibiotic resistant pathogens. Using fecal matter-derived microbiomes we identified a specific community that selected for a carbapenem-resistant *Klebsiella pneumoniae* strain. This selective advantage was due to mutations arising in a LacI-type transcriptional regulator, GlyR, which upregulated expression of the downstream glycoporin GlyP, causing the effect. By deconvoluting the microbiome composition, we identified a focal *E. coli* strain as a central driver of the selection, which was further modulated by other microbiota members. We further demonstrate that the selective advantage was due to carbohydrate competition, and in particular for glycerol-containing compounds. Importantly, *glyR* mutations are under strong positive but conditional selection in clinical *K. pneumoniae* isolates. This implies a reduced competitiveness in other environments, which we experimentally validated *in vitro*. Overall, this study offers a path to identify microbiome-specific interactions that modulate the competitiveness of antibiotic resistant pathogens.

## Introduction

The spread of antibiotic-resistant pathogens, combined with the lack of development of novel antibiotics, is a major challenge to modern medicine. In addition to reduced capacities to treat life-threatening infections, the spread of resistance jeopardizes life-saving medical procedures that rely on prophylactic antibiotic treatment, such as chemotherapy or surgical procedures^1^. In 2019 alone approximately five million deaths were associated with antibiotic resistant pathogens^2^. As the spread of resistant pathogens increases, deaths linked to them are expected to increase significantly over the next two decades if no decisive action is taken^3^. Despite the immense health and economic burden imposed by antibiotic resistance, development of novel antibiotics has lagged behind in the decades following the ‘golden era of antibiotics’ between 1940 to 1960, when most major classes of antibiotics were discovered^4,5^. Many reasons have contributed to this, including both challenges in identifying newly acting compounds and the broken antibiotic market. The result is that hardly any novel antibiotic classes have been introduced in clinical use for decades^4^, with some hope that the tide will change with a new antibiotic class entering the market recently^6^ and more candidates being in the pipeline^7–9^. Instead, most approved compounds in the last decades have been derivatives of antibiotics already in clinical use and are therefore prone to existing resistance mechanisms^10^. In addition to new antibiotics, the improvement of existing therapies can also help in dealing with the antibiotic crisis. This can be facilitated by exploiting drug synergies^11,12^, optimizing drug dosing and delivery^13^ or developing inhibitors for resistance determinants^14^.

A different path to deal with the antibiotic crisis is to understand how cells develop resistance to them, and find ways to prevent, delay or counter-select this process. For example, many resistance determinants have been shown to cause cross-resistance (or collateral sensitivity) where increase in resistance to antibiotic A leads to increase in resistance (or susceptibility) to antibiotic B^15–18^. Treatment regimens can therefore be tailored to avoid pharmaceuticals that share resistance patterns or to combine drug pairs that lead to collateral sensitivity^19^. Although our current knowledge of drug collateral sensitivity is limited, especially since it depends on the pathogen and resistance mechanism, recent developments have opened ways to systematically identify, quantify and understand such drug relationships^20^.

In addition to collateral sensitivity to specific compounds, antibiotic resistance acquisition can come with a ‘fitness cost’ in different environmental conditions^21^. Knowledge of such conditions opens the door for strategies to counter-select the resistant strain and/or promote its displacement by a susceptible strain^22^, which can be treated with the antibiotic. Such strategies are particularly relevant for opportunistic pathogens, as removal of resistance can happen at their benign state, allowing for treatment when/if the bacterium becomes pathogenic. Traditionally, such fitness costs of antibiotic-resistance have been measured under axenic lab conditions by quantifying exponential growth rates or maximum yield. While this approach provides several benefits (high reproducibility, easy automation and low costs) and has been successfully employed for mechanistic studies of the effects of resistance acquisition on cellular physiology^23^, it does not capture the complexity of the natural environments where pathogenic bacteria reside and antibiotic intervention takes place. For example, many opportunistic human pathogens reside together with other microbes in our body, where they compete for nutrients and niches^24^. The most prominent such microbial community is the human gut microbiome, which serves both as a reservoir for opportunistic pathogens and antibiotic resistance determinants^25–27^. With its diversity, complexity and multitude of interactions (e.g. from nutritional competition to cross feeding^28,29^), it poses a versatile challenge for the competitiveness of an antibiotic resistant bacterium, and at the same time an opportunity to identify strains, molecules and/or niches that expose the Achilles heel of resistance elements. Yet methods to do this are currently lacking.

Here, we present a method to capture fitness costs of antibiotic resistance within complex gut microbiomes, and use it to characterize the impact of diverse microbiome compositions on the fitness of the human pathogen *Klebsiella pneumoniae* carrying resistance determinants that are pervasive amongst clinical isolates causing hard-to-treat infections worldwide. We identify a specific microbiota that selects for the antibiotic-resistant isolate to evolve mutations in a LacI-type transcriptional regulator, which gives the strain a competitive advantage over its susceptible parent in utilizing glycerol containing sugars. This mutational signature emerges recurrently in clinical isolates, suggesting that the observed selective pressures exist in nature. At the same time, such variants seem to be an evolutionary dead end in clinics, likely due to reduced competitiveness in other environments. Consistently we show that these microbiome dependent selections are accompanied by severe tradeoffs in utilization of other carbohydrates, which can be used for their rapid counter-selection. Overall, this study exemplifies how we can utilize fecal matter derived microbiomes to identify (counter-)selective pressures for resistant pathogens, which are prevalent *in vivo*.

## Results

### A high-throughput method to quantify resistance-associated fitness costs within gut microbiomes

Bacterial fitness is typically determined by growth curve assays in axenic conditions or by competition assays where the change in ratio of two competing strains is determined over several passages^30^. The pairwise competition assays are more sensitive and can detect interaction-dependent selective effects that are missed in monocultures. We decided to adapt the classical pairwise-competition assay so it could be performed within a complex microbial community setting, and assess how diverse microbiomes impact the competition of the two focal strains (Fig. 1a, b; Methods).

**Figure 1.**
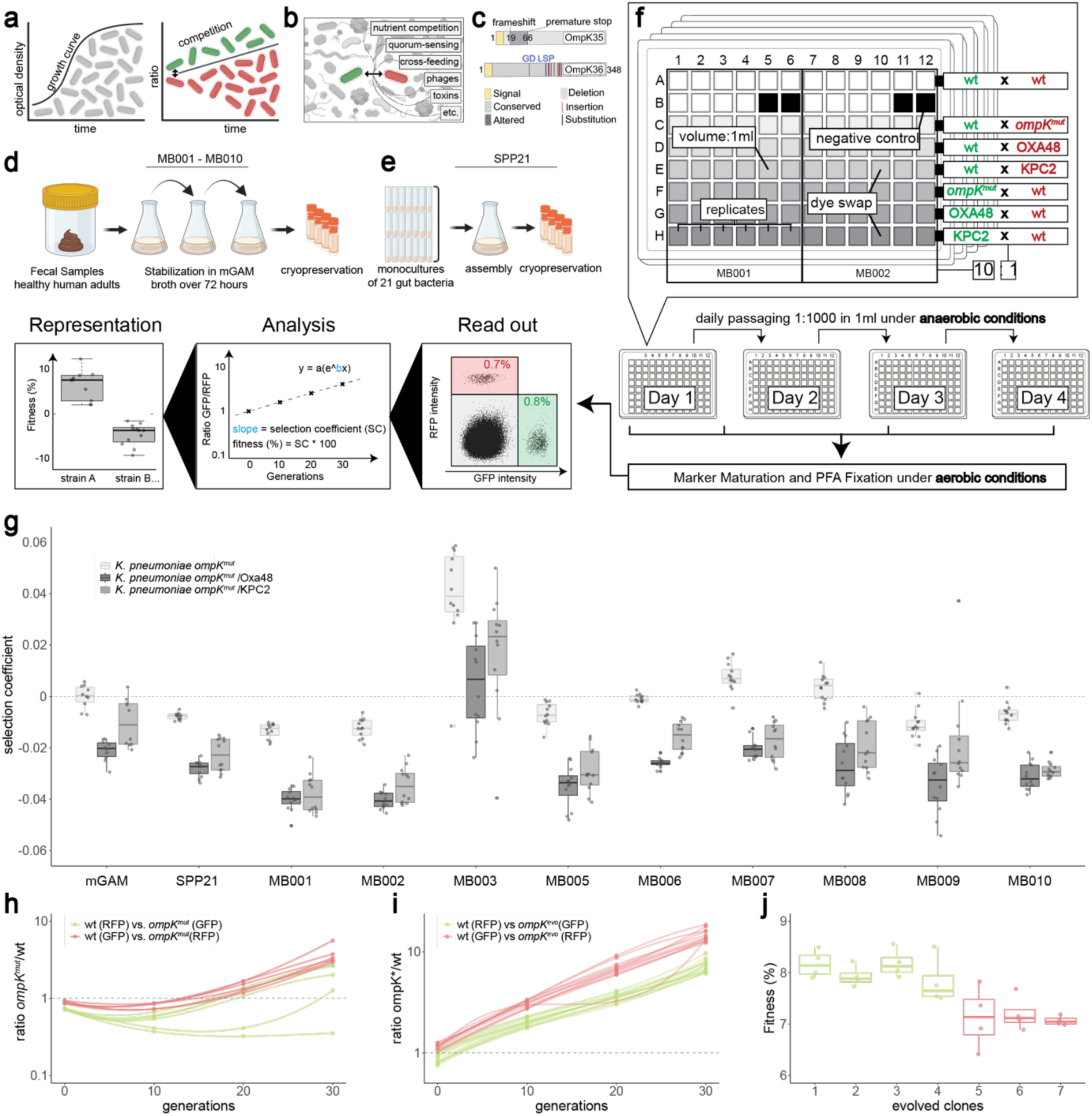
A sensitive method for identifying specific microbiotas that (counter-)select for an antibiotic resistant *K. pneumoniae* mutant. a, Conventional methods to determine bacterial fitness *in vitro*. Fitness is measured by determining lag time, exponential growth rate and maximum yield of an axenically grown culture, or by competing two differently tagged isogenic strains over time to determine their change in ratios, and thereby the selection coefficient. **b,** Schematic representation of factors influencing bacterial fitness in complex communities. **c,** Mutations present in the *K. pneumoniae* porin mutant. *ompK35* carries a frameshift mutation resulting in a premature stop after 66 amino acids causing a complete loss of function. *ompK*36 carries a series of clinically prevalent mutations including a GD-insertion resulting in a restriction of the pore diameter. **d,** Preparation of cryopreserved stool- derived microbial communities MB001 to MB010. Fecal samples of healthy human adults were mixed 1:1 with PBS and diluted in mGAM broth, passaged anaerobically for 3 days in fresh medium and cryopreserved in presence of 12.5% glycerol. **e,** Preparation of cryopreserved synthetic community SPP21. Overnight cultures of 21 prevalent gut bacterial species mixed at equimolar ratios. The synthetic community was grown overnight and cryopreserved in presence of 12.5% glycerol. **f,** Experimental setup of the competition experiment between various antibiotic-resistant *K. pneumoniae* strains in presence of different human gut microbiomes. Both stool-derived and synthetic communities were passaged twice after reviving from glycerol and before competition started. Competitions were performed in 96 deep-well plates with daily passing over 3-4 days under anaerobic conditions with the indicated experimental parameters. Maturation of the fluorescent proteins, PFA-fixation and readout using flow cytometry were performed aerobically. **g,** Selection coefficients of *K. pneumoniae ompK35/ompK36* mutant (annotated as *K. pneumoniae ompK^mut^*) with or without OXA48 or KPC-2 compared to an isogenic wildtype strain in absence (mGAM) or presence of 10 different fecal or synthetic microbiomes. All competitions were performed in a minimum of 5 replicates per marker combination, and all resistant mutant were competed against the wild type with GFP and RFP (with the mutant carrying the corresponding complementary marker). The GFP or RFP associated fitness cost was determined in a wildtype versus wildtype competition and used for subsequent correction of selection coefficients (see methods). Center lines show the medians; box limits indicate the 25th and 75th percentiles; whiskers extend 1.5 times the interquartile range from the 25th and 75th percentiles, data points are plotted as dots. **h,** Change in ratio of *ompK^mut^* over *K. pneumoniae* wildtype in presence of MB003 over 4 days. Colors indicate the marker carried by the porin mutant. **i,** Change in ratio of *K. pneumoniae ompK^evo^* over the wildtype in presence of MB003 over 4 days. Colors indicate if the evolved mutant was isolated from the GFP- or RFP-tagged lineage of *ompK^mut^* j, Selection coefficient of seven *K. pneumoniae ompK^evo^* endpoint clones over the wildtype in presence of MB003. The selection coefficients were corrected for the marker cost determined by the wildtype-wildtype competition in presence of MB003. Colors as in panel **i**.

We chose *K. pneumoniae* as our model strain due to its classification as a priority human pathogen^2,31^, which is frequently associated with antibiotic resistance in clinical settings^32^. *K. pneumoniae* is also a common member of the human gut microbiome^33^, well studied, easy to cultivate and genetically manipulate. We used a previously characterized isogenic set of *K. pneumonia*e strains, which were chromosomally tagged with GFP or RFP markers to allow for sensitive quantification with flow cytometry, and carried antibiotic resistance determinants with minimal fitness effects during axenic growth in nutrient rich conditions^34^. These comprised chromosomal mutations resulting in a loss-of-function deletion and pore-size restriction of the two major porins OmpK35 and OmpK36, respectively (hereafter termed *K. pneumoniae ompK^mut^*) (Fig. 1c), and two clinically prevalent antibiotic resistance plasmids encoding the carbapenemases KPC-2 and OXA-48 (pKpQIL-KPC2 and pOXA-48a). We set out to perform pairwise competition assays between different versions of the resistant *K. pneumonia*e strain and its susceptible parent, each carrying a different fluorescent marker (and the marker swap; see methods) in different complex microbiota communities.

To assess different microbial community compositions, we chose nine native microbiomes derived from fecal samples of healthy human adults (Fig. 1d), and one model synthetic community of 21 different bacterial species that are prevalent in the human gastrointestinal tract^35^ and can provide colonization resistance to pathogens^36^ (Fig. 1e). All communities were stabilized in mGAM broth under anaerobic conditions for 72 hours with daily passaging, before the two *K. pneumoniae* competitors were spiked into the different microbiomes at a 1:1:20 ratio and the community mix was passaged anaerobically for 4 days at a 1:1000 dilution every 24 hours, allowing for 10 doublings per day on average. After each passage, ratios of the competitors (GFP or RFP positive), and the overall microbiome (untagged) counts, were determined using flow cytometry. Thereby we could calculate the selection coefficients of the competitors, i.e. the fitness effect per cell per generation, as well as the carrying capacity of the different microbiomes for *K. pneumoniae*. Adapting the experimental setup to a 96 well format, and using liquid-handling robots and a plate-based spectral analyzer, allowed us to scale up the number of mutants/microbiomes tested, while maintaining high sensitivity (Fig. 1f). The initial total relative abundance of 10% *K. pneumoniae* when the competition coculture was assembled decreased in all conditions by varying magnitudes. Most microbiomes showed a steady decrease across every passage, suggesting that *K. pneumoniae* cannot easily engraft in some of these communities under the nutritional conditions present in mGAM. Abundance stabilized at approximately 3-4% for the microbiomes, MB003 MB005 and MB007, and to < 1% for MB009 (Extended Data Fig. 1).

In the absence of a microbial community, *K. pneumoniae ompK^mut^* was as fit as the wildtype, whereas acquisition of the pKpQIL-KPC2 and the pOXA-48a plasmid led to a minor fitness cost of roughly 1 and 2%, respectively (Fig. 1g). Most microbiomes did not cause a significant change in the competitiveness of the antibiotic-resistant mutants or led to a slight reduction in the fitness of *K. pneumoniae ompK^mut^* and of its derivatives carrying the resistance plasmids – strongest effect being 1.3-2% for MB001. In contrast to all other microbiomes, MB003 led to *K. pneumoniae ompK^mut^* outcompeting its parental susceptible wildtype by about 4% per cell per generation – at least during later passages (Extended Data Fig. 2). The effect was ameliorated by the presence of pKpQIL-KPC2 and pOXA-48a, as those plasmids conferred fitness costs already in the absence of community. In contrast to all other communities, the plasmid carrying variants were not outcompeted by the wildtype in presence of MB003 (Fig. 1g). Altogether these results pointed to antibiotic resistance determinants having microbiome-dependent fitness costs, and highlighted a specific effect of MB003 that led to selection of the antibiotic resistant *K. pneumoniae* porin mutant.

### Microbiome MB003 drives the evolution of a high-fitness variant of the *K. pneumoniae* porin mutant

The ratio changes in the competition experiments were constant over time across most conditions, indicating stable fitness differences of the competitors throughout the experiment (Extended Data Fig. 2 and 3), and allowing us to calculate selection coefficients (Fig. 1g). However, replicates of the three pairwise competitions (*K. pneumoniae ompK^mut^*, *K. pneumoniae ompK^mut^* /pOXA48 and *K. pneumoniae ompK^mut^* /pKPC-2 versus *K. pneumoniae* wild type) showed a large variation in MB003 (Fig. 1g). In fact, *K. pneumoniae ompK^mut^* was initially outcompeted by the susceptible wildtype in presence of MB003, and only increased in ratio over the wildtype at later passages (Fig. 1h). This trajectory change during competition experiments is indicative of a secondary mutation being selected during the experiment time course that provides a fitness advantage to one of the competitors. To validate that an underlying genetic change was driving the competitive advantage of the porin mutant in MB003, we isolated the *K. pneumoniae ompK^mut^* clones from the last day of the competition experiment (hereafter termed *K. pneumoniae ompK^evo^*), and competed them against the wildtype. In line with our hypothesis, we observed a strong and consistent selective advantage of *K. pneumoniae ompK^evo^* (Fig. 1i) resulting in a fitness advantage of 7-8% per cell per generation over the wildtype (Fig. 1j). The fitness advantage was specific for MB003, as the evolved clones did not outcompete the wildtype in presence or absence of any of the other microbiomes – with the exception of MB008 which resulted in a moderate fitness advantage of 2% per cell per generation over the wildtype (Extended Data Fig. 4). Altogether we show that MB003 provides a distinct selective milieu that drives the evolution of a *K. pneumoniae ompK^mut^* high-fitness variant able to outcompete the parental susceptible wildtype.

### A resident *E. coli* strain is driving the selection of a *K. pneumoniae* high-fitness resistant mutant in microbiome MB003

To determine the causative MB003 community member(s) for the enrichment of *K. pneumoniae ompK^evo^* we performed a metagenomic analysis of all nine fecal-derived microbiomes (Fig. 2a). All microbiomes exhibited similar Shannon diversities, and contained between 200 and 450 molecular operational taxonomic units (mOTUs)^37^, with microbiome MB007 having the least and MB001 having the most mOTUs. Strikingly, MB003 which strongly enriched for *K. pneumoniae ompK^evo^*, showed the most distinct metagenomic profile with a dominance of Prevotellaceae. To identify which member(s) of microbiome MB003 contributed to the selection of the high-fitness *K. pneumoniae ompK^evo^* mutant, we chemically perturbed MB003 in order to generate 48 subcommunities with reduced microbial complexity and differential species abundances, as measured by 16S rRNA sequencing (see methods). All subcommunities were then diluted in mGAM and inoculated with the *K. pneumoniae* competitor pair, consisting of *ompK^wt^* and the high-fitness variant *ompK^evo^*. To exclude, that the different conditions used to enrich the subcommunities affect the fitness of the different *K. pneumoniae* strains, we performed the competition in pure mGAM. In contrast to previous competition experiments we performed the co-culture competition experiment only for one day as prolonged passaging in mGAM (without the subcommunity selective condition) would likely shift the subcommunity composition causing inconsistent selective conditions. While this experimental setup is less sensitive than the previous competition experiments and does not allow us to calculate selection coefficients, we could qualitatively screen all subcommunities for their selective effect on the evolved porin mutant by determining a change in ratio from the 1:1 starting mixture to that after 24h of growth in presence of different subcommunities (Fig. 2b). Most subcommunities enriched *K. pneumoniae ompK^evo^* over the wildtype in 24h of growth, but several subcommunities did no longer exhibit this selective effect. The abundance of several genera showed a positive correlation with selection of *K. pneumoniae ompK^evo^* over the wildtype, such as *Sutterella, Parabacteroides* and *Escherichia* (Extended Data Fig. 5). The latter was of particular interest, as certain conditions, such as bile salts, resulted in sub-communities dominated by *E. coli,* while maintaining their selective effect on *K. pneumoniae ompK^evo^* (Fig. 2b). To test whether *E. coli* is sufficient to drive the selection of *K. pneumoniae ompK^evo^*, we isolated the resident *E. coli* s from MB003 (hereafter called *E. coli*^MB003^) and showed that co-culturing it with the *K. pneumoniae* competitors was enough for the *K. pneumoniae ompK^evo^* to outcompete its susceptible wildtype by about 2% (Extended Data Fig. 6). Although subcommunities depleted for *E. coli* (such as after sparfloxacin or cefotetan treatment) were still selective for *K. pneumoniae ompK^evo^* (Fig. 2b), we show that *E. coli* is sufficient to drive part of this selection.

**Figure 2.**
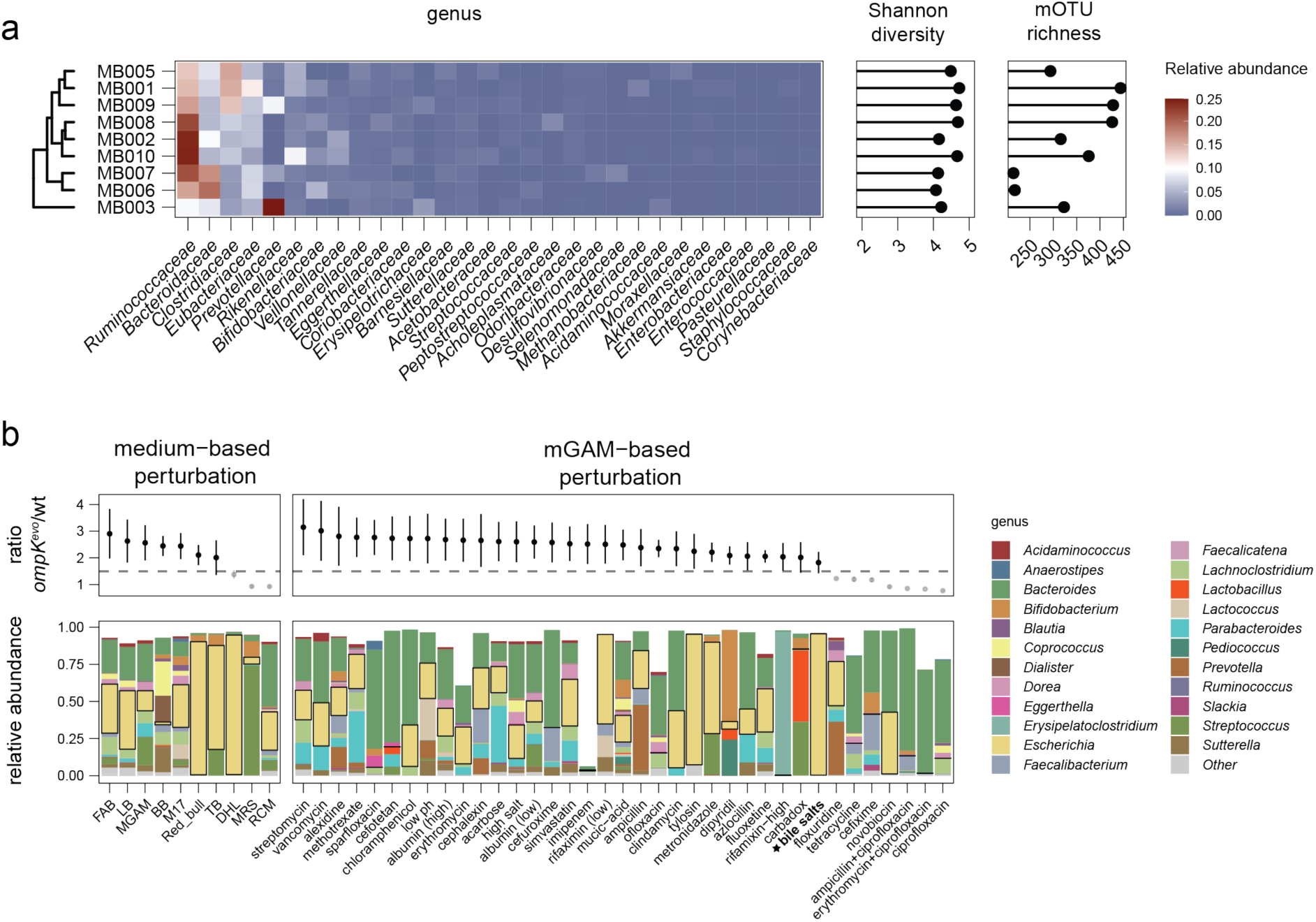
Identification of Microbiome MB003 community members driving the enrichment of *K. pneumoniae ompK^evo^*. a, Composition, Shannon diversity and mOTU richness of the fecal derived communities MB001-MB010 based on metagenomic analysis (see methods). **b,** Ratio of *K. pneumoniae ompK^evo^* over the wildtype after one day of co-culture in different MB003- derived subcommunities. The subcommunities were generated by daily passaging of MB003 under various selective conditions for 3 days. The data points in the upper panel (n = 4 replicates, error bars represent standard deviation) correspond to relative abundance of *K. pneumoniae ompK^evo^* over the wildtype after passaging a 1:1 mixture in the subcommunities for 24hours (secondary y axis). The lower panel shows the microbial composition based on 16S rRNA sequencing. *Escherichia* abundance is accentuated with a black outline.

**Figure 3.**
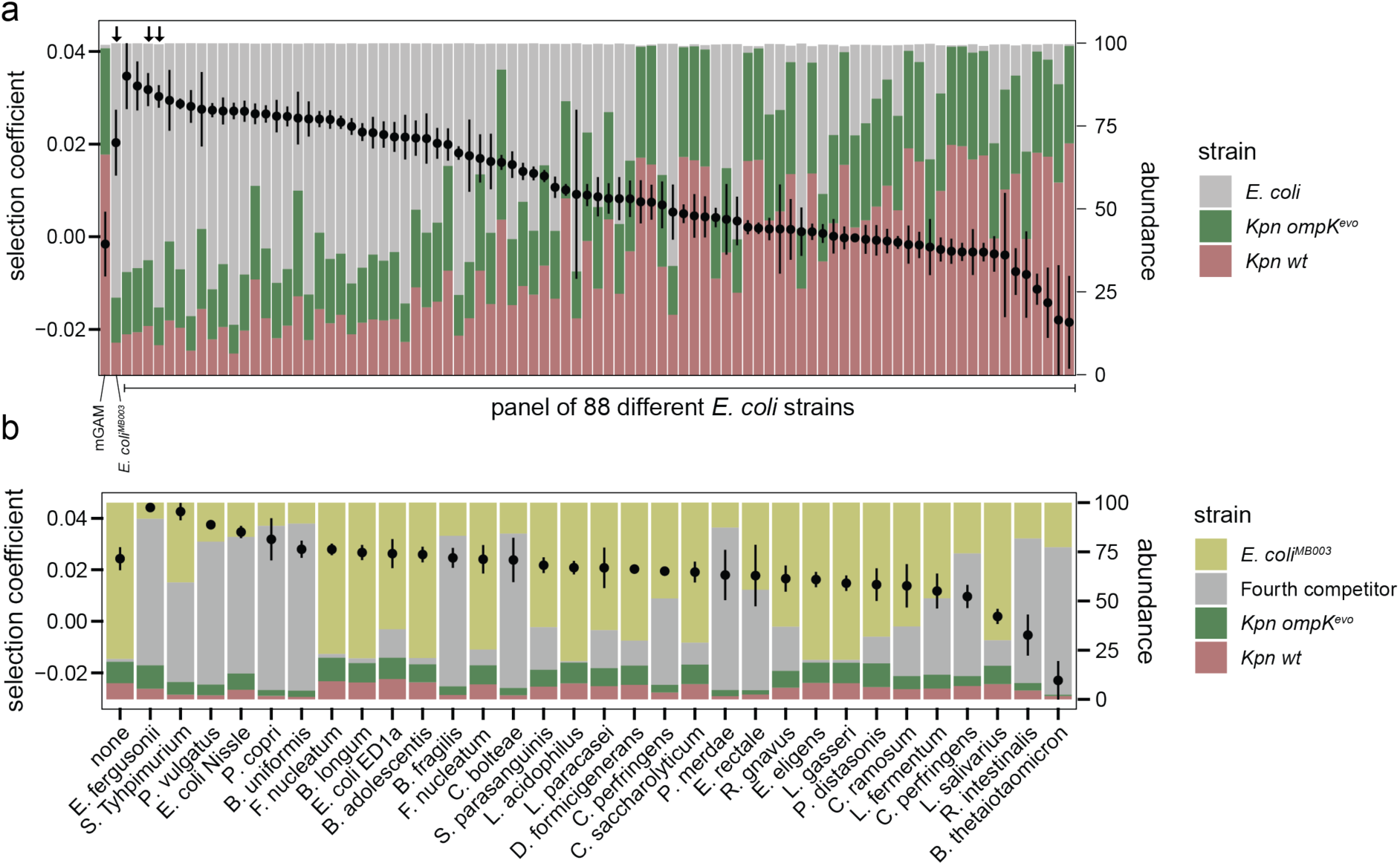
The selective advantage of *K. pneumoniae ompK^evo^* is driven by many *E. coli* strains and depends on higher-order interactions with other gut bacteria. a, Selective advantage of *K. pneumoniae ompK^evo^* over the wildtype in the presence of different *E. coli* strains. The competitions were performed with each *E. coli* strain individually (triple competitions). Data points (primary y-axis) represent the average selection coefficient of *K. pneumoniae ompK^evo^* over the wildtype of four replicates. Error bars represent the standard deviation. The stacked bars (secondary y-axis) represent abundance of the indicated strains at the end point of the competition experiment as determined by flow cytometry. Arrows above bars indicate *E. coli* strains that were isolated from the fecal microbiomes MB003, MB001 and MB005 (left to right). **b,** Selective advantage of *K. pneumoniae ompK^evo^* over the wildtype in presence of YFP- labelled *E. coli^MB003^* and various fourth competitors. The competitions were performed in presence of each fourth competitor individually (quadruple competitions). Data representation as in panel **a**.

### High-order community interactions can modulate the selective advantage *E. coli* provides for *K. pneumoniae ompK^evo^*

*E. coli* is a common member of the human gut microbiome^38^ and present in all fecal microbiomes used in this study (Extended Data Fig. 7), raising the question why the selection of *K. pneumoniae ompK^evo^* occurred only in presence of MB003. To determine how specific the observed effect was to the *E. coli* strain isolated from MB003, we performed competitions of *K. pneumoniae ompK^wt^* and *ompK^evo^* in the presence of different *E. coli* strains. These came from a diverse panel of 85 laboratory, environmental and pathogenic isolates^39^, and included the three *E. coli* strains isolated from MB001, MB002 and MB003 (Fig. 3a). The triple competitions resulted into a wide range of selective coefficients for the *K. pneumoniae ompK^evo^*, ranging from no to approximately 4% fitness benefit. Notably in most cases in which there was low/no selection for *K. pneumoniae ompK^evo^*, this was linked to a poor ability of *E. coli* to engraft in the community (Fig. 3a). This suggests that the selective effect exerted by *E. coli*^MB003^ is a more general trait of *E. coli* rather than being specific to the MB003 strain. Corroborating this, other *E. coli* strains isolated from fecal microbiomes used in this study also exhibited strong selection (> 3% per cell per generation), even if their resident microbiome did not select for the evolved porin mutant (Fig. 1g, 3a).

Given that the selective effect is generally provided by many *E. coli* strains, and that MB003 is not enriched in *E. coli* compared to the other microbiomes (Extended Data Fig. 7), it implies that other microbiota members, besides the residential *E. coli* strains, play a role in the selective effect and significantly modulate its magnitude. To address such potential higher- order interactions, we performed quadruple competition experiments including as a fourth competitor one further bacterial species relevant to the human gut microbiota (Fig. 3b). In the competition experiment, we inoculated *K. pneumoniae ompK^evo^*, *K. pneumoniae* wildtype, *E. coli^MB0^*^03^ and the fourth competitor in equal proportions and followed their abundance over three days of competition. We also labeled *E. coli^MB003^* with a chromosomally encoded YFP marker to be able to follow the abundance of all four strains in the competition. The presence of a fourth competitor (untagged) modulated the selective advantage of *K. pneumoniae ompK^evo^* exerted by *E. coli* in different ways. For example, the presence of other enterobacteria or *Prevotella copri* increased the selective advantage of *K. pneumoniae ompK^evo^* in presence of *E. coli^MB003^* to 3 percent, while the presence of *Clostridium perfringens* strongly alleviated the fitness advantage. Notably, these modulating effects align with compositional differences observed in MB003, which exhibits lower abundance for *Clostridiales* and higher abundance of *Prevotella* compared to the other fecal microbiomes (Fig. 2a). Collectively, our results demonstrate that while *E. coli* is sufficient to provide a selective advantage of *K. pneumoniae ompK^evo^* over its ancestral susceptible wildtype, higher order interactions in the community shape the magnitude and even direction of the selection. This is likely the reason we see the selective advantage only in MB003, and not in other microbiomes.

### Microbiome MB003 selects for *glyR* mutations that lead to upregulation of an alternative porin

To elucidate the underlying molecular mechanism that allowed for the evolved porin-mutant to benefit in presence of MB003, we whole-genome sequenced seven *K. pneumoniae ompK^evo^* clones. All strains had acquired a mutation in a LacI-type transcription factor hereafter termed ‘glycoporin regulator’ (*glyR*). The mutational spectrum spanned frameshifts and substitutions causing premature stop codons, internal deletions and single amino acid substitutions namely T60P, I54S and C248W (Fig. 4a). *glyR* is located upstream of three genes encoding a putative permease, hydrolase and porin. This is a common organization of a carbohydrate catabolic operon where the transcription factor regulates expression of the structural genes, which facilitate sugar uptake through the outer- and inner- membranes, followed by its hydrolysis in the cytoplasm. In line with this hypothesis, whole cell proteomes of all seven isolated *K. pneumoniae ompK^evo^* mutants exhibited consistently strong and specific upregulation of all three proteins encoded in the structural genes (Fig. 4b and Extended Data Fig. 8).

**Figure 4.**
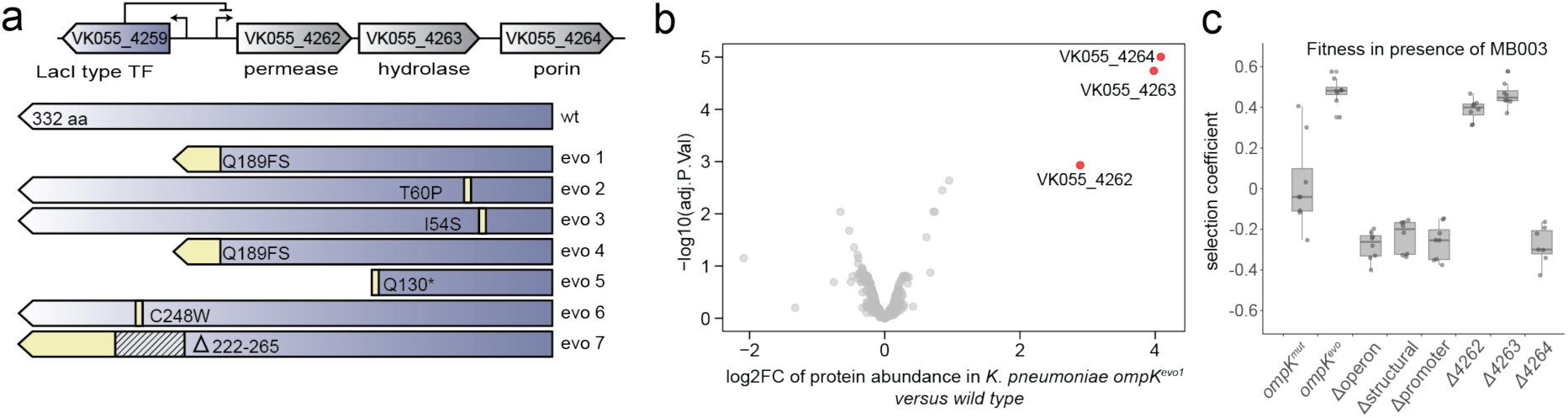
Mutations in a transcription factor in *K. pneumoniae ompK^evo^* cause the fitness advantage in MB003, due to upregulation of the downstream porin. a, Whole genome sequence analysis of seven *K. pneumoniae ompK^evo^* mutants. All clones carry mutations in a LacI-type transcription factor resulting in premature stops, internal deletions or amino acid substitutions. The transcription factor is located upstream of a carbohydrate catabolic operon consisting of a permease, a hydrolase and an additional porin. The labels indicate the locus tags according to reference genome CP009208. **b,** Proteome abundance changes induced by acquisition of a *glyR* mutation in *K. pneumoniae ompK^mut^*. The wildtype and evolved porin mutant carrying a T60P substitution in the LacI-type transcription factor (evo2) were grown for 16 hours in mGAM under anaerobic conditions and subjected to mass spectrometry. All genes of the carbohydrate catabolic operon are upregulated in the *glyR* T60P mutant. **c,** Selective advantage of various deletion mutants of the carbohydrate-catabolic operon in the *K. pneumoniae ompK^evo^* background or in the wildtype background over the parental strain in presence of MB003. Upregulation of the porin (4264) is causal for the observed selective advantage. Box plots are depicted as in Fig. 1g.

To determine which of the structural genes are causing the fitness advantage of the *glyR* mutant background when grown in presence of MB003, we constructed several knockout variants. Removing the whole operon, the predicted transcription factor binding sites or all three structural genes resulted in a complete loss of the fitness advantage phenotype, confirming that indeed the structural genes upregulated in the *glyR* mutant background are causative. Furthermore, knockouts of each of the individual structural genes revealed that the porin alone was sufficient for the fitness advantage (Fig. 4c). The porin (hereafter termed glycoporin, GlyP) is predicted to belong to the raffinose porin family RafY (Extended Data Fig. 9), which contains a larger pore size compared to OmpK35/36 and is involved in the uptake of a range of carbohydrates including oligosaccharides^40,41^.

### Carbohydrate competition drives the selective advantage of the *glyR* mutants

There are several potential mechanisms of how an increased expression of GlyP would provide a fitness advantage in a community context. Since porins are involved in nutrient uptake, it is conceivable that the fitness benefit is based on an increased nutritional competence. However, porins also play a role in the uptake of toxins, export of metabolic waste products or as receptors of phage and conjugative plasmids^42–44^. As such, an outer membrane with different porin composition could render *K. pneumoniae ompK^evo^* able to use a new niche or to be more resistant to toxic products or to negative interactions with other community members. To define the underlying mode of action of the evolved mutants, we grew *E. coli*^MB003^ until saturation (20h), sterile filtered its supernatant, and used the spent medium to determine growth characteristics of each *K. pneumoniae* strain in monocultures. *K. pneumoniae ompK^evo^* grew significantly better than its parental strains and the wildtype in the spent medium (Fig. 5a), demonstrating that the selective effect mediated by *E. coli* is contact independent. Supplementing the spent medium with a good carbon source (0.2% glucose), resulted in higher growth yields and similar growth characteristics of all three *K. pneumoniae* strains (Fig. 5b), indicating that: a) the lack of carbon sources is growth-limiting for *K. pneumoniae* in these conditions, and b) the selection advantage of *ompK^evo^* is only visible in poor nutrient conditions and hence most likely works via increased nutritional competence, as growth inhibitory mechanisms like toxin uptake or phage susceptibility would have manifested independently of the absence or presence of glucose.

**Figure 5.**
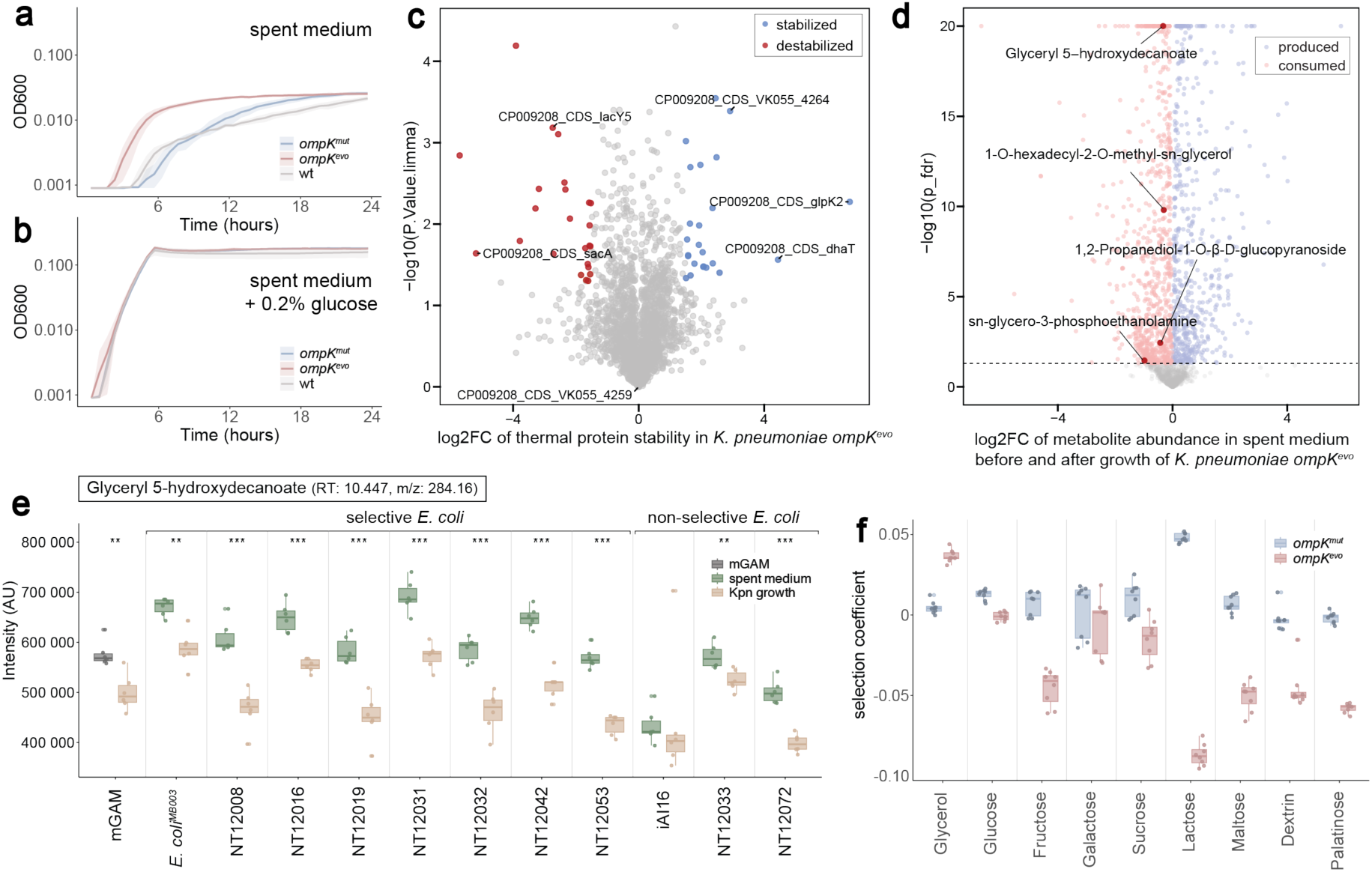
GlyP upregulation allows *K. pneumoniae* to benefit from glycerol containing compounds. a, Growth curves of *K. pneumoniae* wildtype, *ompK^mut^* and *ompK^evo^* in *E. coli^MB003^* spent medium without supplements or **b,** with supplementation of 0.2% glucose. The spent medium was prepared by growing *E. coli^MB003^* overnight in mGAM at 37 °C under anaerobic conditions and subsequent sterile filtration with a 0.22µM pore size filter. Lines represent average of four independent replicates, confidence intervals represent one standard deviation. **c,** Change in thermal stability of the proteome of *K. pneumoniae ompK^evo^* compared to the wildtype in presence of *E. coli^MB003^* spent medium (see methods). Three replicates of *K. pneumoniae ompK^evo^* and *K. pneumoniae* wild type were grown anaerobically overnight in mGAM. Whole-cells lysates were treated with *E. coli^MB003^* spent medium and subjected to thermal proteome profiling. Significant hits are indicated with red (destabilized) and blue (stabilized). Members of the carbohydrate catabolic operon described in Fig. 4a as well as the two strongest stabilized proteins are labelled. **d,** Metabolomics of *E. coli^MB003^* spent medium before and after growth of *K. pneumoniae ompK^evo^*. Compounds significantly enriched after growth of *K. pneumoniae ompK^evo^* are colored in blue, compounds significantly diminished are coloured in red. Glycerol-derivatives are labelled. **e**, Signal intensities for Glyceryl 5- hydroxydecanoate in spent media of seven different *E. coli* isolates and in mGAM before and after growth of *K pneumoniae ompK^evo^.* All experiments were performed in six replicates. Box plots depicted as in Fig. 1g. Statistical significance was assessed using a t-test. * p_adjusted_ < 0.05, ** p_adjusted_ < 0.01, *** p_adjusted_ < 0.001 **f**, Fitness of *K. pneumoniae ompK^mut^* and *K. pneumoniae ompK^evo^* compared to *K. pneumoniae* wildtype grown in different carbon sources. Competitions were performed in M9 minimal media supplemented with defined carbon sources as indicated. Box plots are depicted as in Fig. 1g.

MGAM is a rich medium designed to support the growth of a wide range of gut-derived bacteria, and hence it contains a plethora of potential carbon sources that could create a metabolic niche that selects for *K. pneumoniae ompK^evo^*. To narrow down the spectrum of potential candidate compounds, we performed thermal proteome profiling (TPP)^45,46^. Due to the fact that enzyme-substrate interactions lead to thermal stabilization^47^, TPP has the potential to identify catabolic pathways that are thermally stabilized in *K. pneumoniae ompK^evo^*, but not in the *K. pneumoniae* wildtype, in the presence of the *E. coli^MB003^* spent medium. Indeed, by treating whole cell lysates of *K. pneumoniae* wildtype and *K. pneumoniae ompK^evo^* mutant with *E. coli^MB003^* spent medium, we identified several proteins that were thermally stabilized only in the *K. pneumoniae ompK^evo^* lysate (Fig. 5c). In addition to GlyP itself, which was higher expressed in the *ompK^evo^* cells (Fig. 4b), two of the most strongly stabilized proteins in spent-medium treated *K. pneumoniae ompK^evo^* were DhaT and GlpK, both of which are involved in glycerol catabolism. DhaT is the 1,3-propanediol dehydrogenase, which converts 3-hydroxypropionaldehylde (generated from glycerol by a dehydratase) to 1,3- propanediol, and GlpK is the glycerol kinase. The strong stabilization of both enzymes suggested that glycerol, or glycerol-derivatives, were present in the spent media, and that glycerol catabolic enzymes were active in *K. pneumoniae ompK^evo^* but not in the wildtype.

To determine whether glycerol or glycerol derivatives were available in the spent medium, we decided to measure the metabolites present in the *E. coli^MB003^* spent medium before and after growth of *K. pneumoniae ompK^evo^*. We chose eight different *E. coli* strains that were selective for *K. pneumoniae ompK^evo^* as well as three *E. coli* strains that were not (Fig. 3a), and prepared spent medium of each strain individually. By profiling the growth of *K. pneumoniae* wildtype, *K. pneumoniae ompK^mut^* and *K. pneumoniae ompK^evo^* in these spent media, we could confirm that the growth characteristics in the various spent media (Extended Data Fig. 10) were consistent with the selective coefficients determined in the co-culture experiments (Fig. 3a). *K. pneumoniae ompK^evo^* showed a growth advantage in spent media prepared from ‘selective’ *E. coli* strains, while it grew equally well to the wildtype in spent medium prepared from ‘non- selective’ *E. coli* strains (Extended Data Fig. 10). We then subjected all spent media before and after growth of *K. pneumoniae ompK^evo^* to untargeted metabolomics to determine which compounds were produced or consumed by *K. pneumoniae* (Fig. 5d). While we did not detect free glycerol in the samples, we identified several glycerol-containing compounds that were (i) present in mGAM, (ii) not consumed by *E. coli*, and (iii) consumed after growth of *K. pneumoniae ompK^evo^* (Fig. 5e and Extended Data Fig. 11). As such, Glyceryl-5- hydroxydecanoate was significantly reduced in abundance in all spent-media prepared from ‘selective’ *E. coli* isolates. While also found in spent media derived from non-selective *E. coli*, it was either consumed to a low degree (iAI16 and NT12033) or present at a lower levels (iAI16 NT12072) compared to spent media derived from ‘selective’ *E. coli*. Judging from the robust growth of all *K. pneumoniae* strains in the spent media of non-selective *E. coli* strains (Extended Data Fig. 10), it is likely that other better carbon sources are available to them, akin to when glucose is present in the spent medium of *E. coli^MB003^* (Fig. 5b).

To unambiguously show that selection of *K. pneumoniae ompK^evo^* is due to an advantage to catabolize glycerol, we performed competition experiments of *K. pneumoniae* wildtype vs either *K. pneumoniae ompK^mut^* or *ompK^evo^* in minimal medium supplemented with glycerol or glucose as the sole carbon sources. While *K. pneumoniae ompK^mut^* grew similarly to the wildtype in glycerol, the *K. pneumoniae ompK^evo^* exhibited a significantly increased fitness of around 4% per cell per generation compared to the wildtype. This fitness advantage was specific to glycerol, as the evolved mutant grew similarly as the wildtype with glucose as the sole carbon source (Fig. 5f). To determine if the selective advantage of *glyR* mutants is specific for growth on glycerol, we tested a series of additional carbon sources for their impact on fitness of *K. pneumoniae ompK^mut^* and *K. pneumoniae ompK^evo^* compared to the wildtype. In stark contrast to growth on glycerol, we observed that other carbon sources, such as fructose, lactose or maltose were disfavoring the *K. pneumoniae ompK^evo^* with counterselection coefficients reaching up to 9% per cell per generation.

In summary our results show, that treatment of *glyR* mutant lysates with *E. coli* spent medium causes thermal stabilization of glycerol catabolic enzymes (indicating their metabolic activity), that mGAM contains glycerol-derivatives that are consumed by *K. pneumoniae*, but not by *E. coli*, and that the *glyR* mutant has a selective advantage when grown in presence of glycerol as the sole carbon source. This suggests that specific microbiota composition(s) can create a niche for consumption of glycerol derivatives. This drives the selection of mutations in *glyR*, which lead to increase in GlyP expression in the *ompK35/36* mutant.

#### *glyR* mutations are under strong positive selection in clinical *K. pneumoniae* populations

We next exploited a large collection of publicly-available genomes of the globally-important high-risk lineage, clonal complex (CC) 258, which includes ST258 and other closely-related STs. Thereby we could assess the sequence variation and evolution within the *glyP* operon, and determine if the selective pressures identified in our *in vitro* screen are clinically relevant. In particular, we generated a mapping-based chromosomal alignment of 9214 high-quality CC258 genomes and used an ancestral reconstruction approach that, for each chromosomal base position, inferred the ancestral base state at all nodes across a recombination-free phylogenetic tree. For each base position, we analyzed the occurrence and frequency of base (state) changes across the phylogenetic tree, with a focus on the *glyP* operon. While the structural genes within the operon showed only a few base changes with no more than 3 SNPs occurring at any individual base position, we identified a single mutational hotspot (C->A) at position 4,366,094 (with respect to the HS11286 reference genome), corresponding to the T60P substitution in GlyR (Fig. 6a). We estimate that this substitution occurred independently 35 times across the CC258 phylogeny. Interestingly, we identified the same substitution in our *in vitro* screening (Fig. 4a), confirming that the *in vitro* selection driven by MB003 in mGAM is recapitulating selective pressures acting on clinical *K. pneumoniae* populations. Notably, we also identified a distinct subclade in which *glyR* is entirely absent due to a deletion spanning the gene (Fig. 6b, clade B). Consistent with this pattern, our *in vitro* screen also identified premature stop codons and frameshift mutations in *glyR* (Fig. 4a).

**Figure 6.**
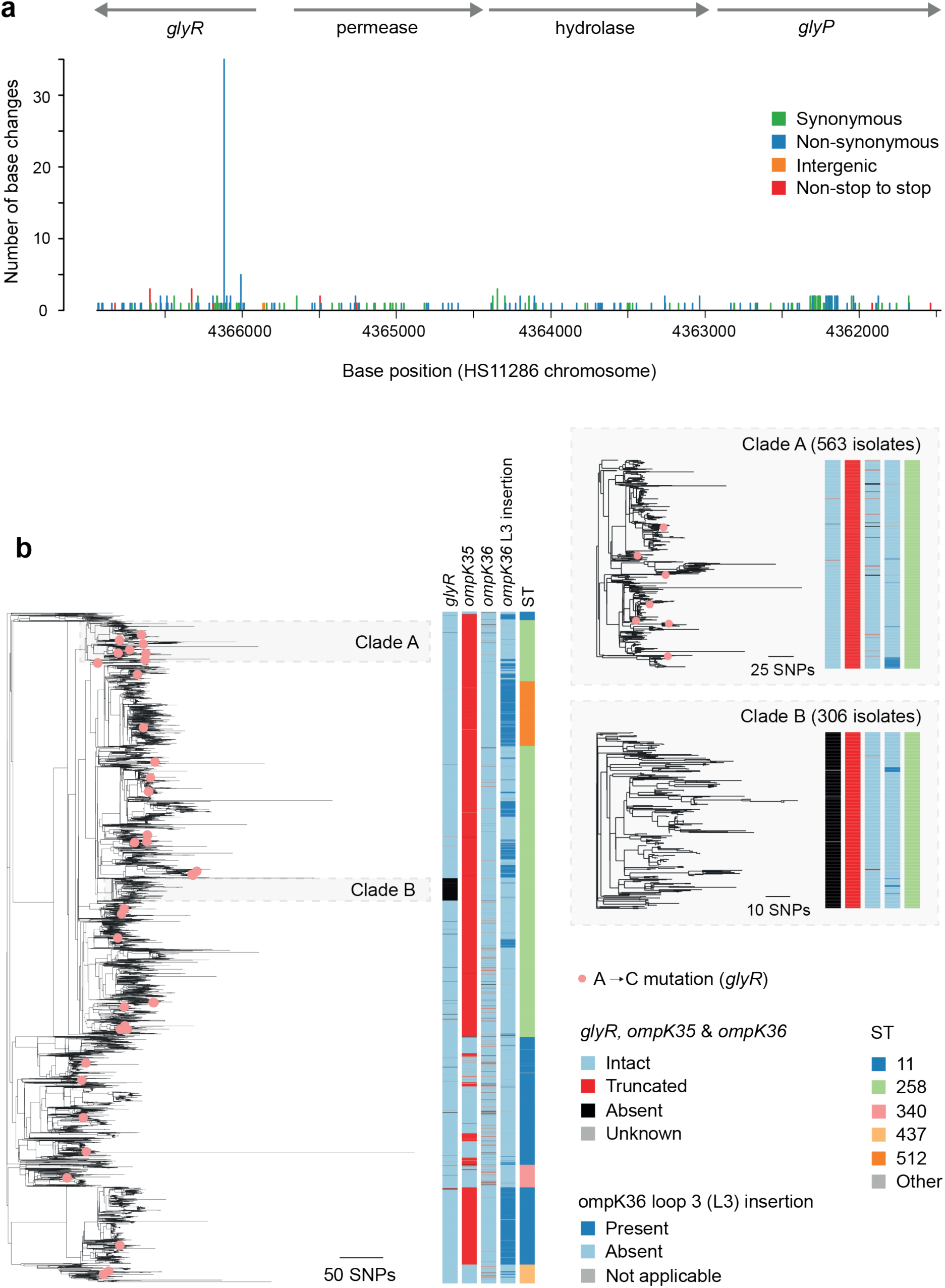
***glyR* T60P substitution is under conditional positive selection in clinical isolates of *K. pneumoniae*. a,** Number of base (state) changes inferred to have occurred across the phylogeny of 9214 CC258 isolates at each base position spanning the length of the *glyR*/*glyP* operons and with respect to the HS11286 reference genome. Base changes are colored by their effect on the protein sequence. **b**, Phylogenetic tree of 9214 CC258 isolates constructed after the removal of recombined regions from the alignment. Isolate nodes are colored pink if they possess a “C” at position 4366094 (with respect to the HS11286 reference), corresponding to amino acid position 60 within the LacI type TF. Metadata columns show whether the *glyR*, *ompK35* and *ompK36* genes are intact, truncated or absent, the presence or absence of an *ompK36* loop 3 insertion (GD or TD) and the ST of the genomes. Clades A and B are shown with additional resolution on the right-hand side. The scale bars represent the number of SNPs. An interactive vizualisation of this tree, together with additional metadata and genotypic data, is available via Microreact at https://microreact.org/project/kp-cc258.

Despite the *glyR* T60P mutation emerging frequently across the CC258 population, its sporadic occurrence on terminal or near-terminal branches of the phylogenetic tree, and subsequent presence in only 41/9214 (0.4%) of CC258 genomes, suggests that this mutation cannot expand clonally (Fig. 6b). This is in contrast to what is expected for mutations that are under long-term positive selection, which would expand within larger clades of the tree. This absence of clusters is indicative, that *glyR* mutations are likely under strong positive selection in some conditions, but then become disadvantageous and are quickly counter-selected when conditions change. This is in line with our findings, that *K. pneumoniae ompK^evo^* is only selected in certain microbiomes (Fig. 1g), and that *K. pneumoniae ompK^evo^* has no benefit, or even strong fitness defects, when growing on other carbon sources than glycerol (Fig. 5f). Together, these findings show that GlyP upregulation bestows a conditional selective advantage for *glyR* mutations in *K. pneumoniae* in the presence of glycerol-containing niches, but these mutations are an evolutionary dead-end when the available niche changes.

## Discussion

Here, we determined the impact of diverse human gut microbiotas on the fitness of a prevalent antibiotic resistance mutation in the ESKAPE pathogen *K. pneumoniae*. We discovered that a specific microbiota drove the evolution of a high-fitness variant in the *K. pneumoniae* resistant strain. The fitness benefit was caused by mutations in the transcriptional regulator *glyR*, which resulted in the upregulation of an alternative porin, GlyP. By using proteomic, metabolomic and phenotypic analyses we identified nutrient competition for glycerol-containing compounds as the driving factor of the selection.

The general exclusion of intruding, usually pathogenic, species by the residential microbiota is termed colonization resistance, and is an important property of a healthy microbiome. Various studies have shown the benefit of resident/probiotic bacterial species in preventing the invasion of pathogenic isolates^48–51^. Carbohydrate competition has been shown to play a key role in colonization resistance against various pathogens, including, *E. coli*^50^, *K. pneumoniae*^49,52^ and *S. enterica* serovar Typhimurium^48,49^, where direct competition for consumption of specific carbon sources such as galactitol or beta-glycosides can restrict growth of pathogenic bacteria. Here we show that nutrients, and specifically glycerol catabolism, drives the evolution and enrichment of an antibiotic resistant *K. pneumoniae* porin-mutant. Although we showed that *E. coli* was at least partially responsible for the creation of this selective niche, higher-order interactions with other community members augmented or removed the selection. For example, *Clostridium perfringens* removed the selective niche. Interestingly, *E. coli* is unable to utilize glycerol as the sole carbon source under strictly anaerobic conditions, while *K. pneumoniae* and members of the *Clostridiales* family can^53–55^. Thus, we propose a model where *E. coli* is exerting its selective effect for the high-fitness K*. pneumoniae* porin-mutant not by direct carbohydrate competition, but rather by creating a niche by removing other, more favorable carbon sources.

Furthermore, we show that the selection for the high-fitness *K. pneumoniae* porin mutant is context-dependent, as the *glyR* mutations are detrimental in other conditions. Clinical *K. pneumoniae* isolates seem to be subjected to similar selective pressures, as shown by the recurrent emergence of the *glyR* T60P mutation among clinical isolates. At the same time, and in line with the results we presented here, the selective pressures are highly conditional, limiting the clonal expansion *of K. pneumoniae* carrying *glyR* mutations. It is conceivable that these mutations occur in isolated patients with favorable microbiome composition and dietary habits, but are not competitive when the environment changes (diet or microbiome) restricting their expansion and transmission. Identifying and understanding selective pressures against (high-fitness) antibiotic-resistant pathogens could enable us to drive off antibiotic resistant strains by dietary modulation.

Strategies for decolonization of pathogens from microbiomes are currently being discussed and actively researched^56^. However counter-selection of an opportunistic pathogen from a ‘healthy’ microbiome opens a niche that could be covered by other invading pathogens. In contrast, finding conditions that specifically counter-select antibiotic-resistant variants of native opportunistic pathogens of the human gut microbiome (e.g. in *K. pneumoniae* or *E. coli*) would avoid this opening of the niche for new invaders. Instead this would allow the antibiotic susceptible strains of the same species to replace their resistant counterparts, and make treatment easier would the opportunistic pathogen lead to infection. Our study lays out a roadmap for how to identify potential probiotic bacteria and/or dietary compounds restricting growth of antibiotic-resistant bacteria that could subsequently be used to drive out antibiotic resistant pathogens from healthy gut microbiomes, decreasing the risk of acquiring hard-to-treat infections in the future.

## Materials and Methods

### Isolation of fecal microbiome samples

Fresh fecal samples from nine healthy donors were collected and transferred directly into an anaerobic chamber. Samples were mixed at a 1:1 ratio (w/v) with PBS containing 40% glycerol and 0.5 g/L cysteine, homogenized, and aliquoted into cryovials for preservation. At the time of sample collection, one aliquot was inoculated into 50 ml of mGAM medium, serially diluted, and incubated anaerobically at 37 °C for 24 h. Following incubation, 800 µl of the diluted cultures was combined with 200 µl of 50% glycerol and preserved at −80 °C. The collection of human stool samples and all related experiments were approved by the EMBL Bioethics Internal Advisory Committee. Informed consent was obtained from all donors (EMBL BIAC2015-009). The synthetic community was assembled as previously described^35,57^. In brief, monocultures of each species were grown overnight in mGAM broth anaerobically at 37 °C, adjusted to OD_600_ = 0.01, mixed at equimolar ratios and cryopreserved in 15% glycerol. For community experiments, the frozen community was reconstituted in fresh mGAM by a 1:100 dilution and grown for 24 hours in mGAM broth anaerobically at 37 °C.

### DNA extraction

Genomic DNA was isolated in 96-well deep-well plates. Cells were washed in PBS, and pellets were resuspended in cell suspension solution (281 µl) from the GNOME DNA isolation kit (MP Biomedicals). Resuspended cells were enzymatically digested with lysozyme (25 µl; 400,000 U ml−1, Sigma) at 37 °C for 1h. Lysis was further enhanced by three freeze-thaw cycles in liquid nitrogen, followed by the addition of the kit’s lysis solution (15.2 µl) and RNA mix (20 µl). A final lysis step was conducted using glass beads (Glasperlen, Edmund Bühler) with bead- beating (5 min x 2 at 30 Hz) in a Tissue Lyzer II (Qiagen). Lysates were then incubated at 37 °C with shaking for 30 min, followed by protease treatment (12.8 µl; GNOMETM kit) at 55 °C for 2h. Cell debris was removed by centrifugation (3200 g, 5 min) and 200 µl of the supernatant was mixed with 100 μl of TENP buffer (50 mM Tris-HCl pH 8, 20 mM EDTA, 100 mM NaCl, 1% polyvinylpolypyrrolidone). The mixture was incubated for 10 min at 4 °C with 75 µl of salt out solution (GNOME kit), followed by 10 min centrifugation at 3200 g. 200 µl of supernatant was transferred to a fresh plate, mixed with 500 µl of ice-cold ethanol and 70 µl of 3 M NaOAc pH 5.2 were added, and kept at −30 °C overnight to precipitate DNA. The next day, DNA was pelleted (3200 g, 45 min, 4 °C), washed with 70% ethanol, air-dried, and rehydrated in molecular-grade water at 4 °C.

### 16S sequencing

For sequencing, 16S rRNA gene libraries were prepared using a two-step PCR approach with Phire Hot Start II DNA Polymerase (Thermo Fisher Scientific). The V4 region was first amplified, and a second PCR was carried out with indexed primers containing Illumina adaptors. Libraries were sequenced using paired-end chemistry (2 x 250 bp) on a MiSeq platform.

Raw sequencing reads from WGS and 16S data were processed using bbduk (v38.91). For WGS data, reads underwent several preprocessing steps to remove low-quality sequences: Low-quality bases were trimmed from both ends (qtrim = rl, trimq = 3), followed by the removal of entire reads falling below a quality threshold (maq = 25). Adapter sequences were then removed (ktrim = r, k = 23, mink = 11, hdist = 1, tpe = true, tbo = true) and reads shorter than 75 base pairs were discarded. For 16S data, reads underwent low-quality base trimming (qtrim = rl, trimq = 10), followed by targeted removal of 5’ amplicon primers from R1 and any potential reverse complement primers from R2 (ktrim = l, k = 9, mink = 1, hdist = 1, restrictleft = 50, maq = 20). Similarly, 3’ primers were removed from R2, along with any potential reverse complement sequences from R1 (ktrim = r, k = 9, mink = 1, hdist = 1, restrictright = 50, maq = 20). Finally, reads shorter than 75 base pairs were discarded.

### Metagenomic sequencing

Microbial DNA was extracted from collected stool samples using Qiagen AllPrep PowerFecal Pro DNA/RNA Kit (Qiagen, Hilden, Germany) following the manufacturer’s protocol. Metagenomic sequencing libraries were prepared using the NEBNext Ultra II DNA Library Prep kit (New England Biolabs, MA, USA) with a targeted insert size of 350-400bp and Dual Index multiplex oligos. Libraries were prepared using a liquid automated system (Beckman Coulter i7 Series) and sequenced on an Illumina HiSeq 4000 platform (Illumina, San Diego, CA, USA) with 2×150bp paired-end reads.

### Taxonomic profiling

For profiling of 16S data, we used IDTAXA84 from the DECIPHER package (v3.0.0)^58^ with parameters strand = both and threshold = 40 and the GTDB r207 (modified) training set. WGS data were profiled using mOTUs (v3.1)^59^ with default settings.

### Community competition experiments

All competition experiments were performed at 37 °C in an anaerobic chamber (Coy laboratory Products; 2% H2, 12% CO2, 86% N2) in modified Gifu anaerobic medium (mGAM) broth (HyServe, produced by Nissui Pharmaceuticals). For competitions in complex communities, aliquots of the cryopreserved fecal microbiomes were thawed under anaerobic conditions and 50µl were diluted in 5ml MGAM and grown overnight. The subsequent day, the cultures were diluted in fresh mGAM broth to a final OD_600_ of 0.01. Overnight cultures of the fluorescently labeled *K. pneumoniae* competitors were mixed at equimolar ratios and added to the OD_600_-adjusted communities at a final OD600 of 0.001, leaving a 10-fold excess of community members compared to *K. pneumoniae* competitors. Each day, the competition cultures were diluted 1:1000 in fresh mGAM broth to a final volume of 1ml. Each competition experiment was conducted in two configurations: one in which strain A was tagged with GFP and strain B with RFP, and another in which the fluorescent markers were reversed. Each configuration was tested with a minimum of five biological replicates. Additionally, we competed two independently constructed sets of the *K. pneumoniae* wild type carrying either GFP or RFP with 10 biological replicates under each condition (microbiome) to determine the fitness effect imposed by marker expression. The selection coefficients determined for the *K. pneumoniae* wild type tagged with GFP or RFP were subsequently subtracted from selection coefficients of GFP or RFP positive strains accordingly.

Competition in defined communities (triple competitions in presence of *E. coli* or quadruple competition in the presence of an additional fourth competitor) were performed in a similar fashion. Each competitor was grown overnight in mGAM and mixed at equimolar ratios to a starting OD_600_ of 0.01. Competition experiments with defined carbon sources were performed in M9 minimal medium (0.1 mM CaCl_2_, 1mM MgSO_4_, 1x M9 Salts (Sigma Aldrich)) supplemented with the carbon source of interest at a final concentration of 0.2%.

To determine the ratios of GFP^+^, RFP^+^ or GFP^-^/RFP^-^ cells, 50µl aliquots were removed from each passage, washed twice with PBS and samples were resuspended in 50µl PBS. The samples were incubated for 2h at room temperature under aerobiosis to allow maturation of the fluorescent proteins. After two hours cells were fixed by adding an equal amount of 4% Paraformaldehyde and incubating for 10 minutes. After fixation, cells were washed once with PBS and taken up in 150µl final volume PBS. Samples were subjected to flow cytometry to determine the ratio of the RFP^+^ and GFP^+^ competitors, as well as the untagged microbiome using the Aurora spectral analyzer (Cytek Biosciences). In rare cases, individual wells were excluded from analyses where fluorescent populations could not be clearly distinguished from non-fluorescent background events, likely due to incomplete resuspension before aeration causing a failure in fluorescent protein maturation.

### Whole genome sequencing

To isolate the endpoint *K. penumoniae* strains, competition cultures were streaked out on LB (5 g yeast extract, 10 g Tryptone, 10 g NaCl L^-1^) Agar plates containing 50 mg/L Carbenicillin and incubated aerobically at 37°C overnight, which specifically selects for the growth of K. pneumoniae. GFP or RFP tagged colonies were differentiated using a Flu-O-Blu transilluminator (Biozym). Colonies of interest were purified by streaking on LB agar plates and incubating at 37 °C overnight. The isolates were then grown overnight in LB broth and cryopreserved using 15% glycerol. 1 ml of each culture was used to extract genomic DNA using the MasterPure DNA purification kit according to the manufacturer’s recommendations (Epicentre). All samples were subjected to whole-genome sequencing using a MiSeq system (Illumina, USA). The paired-end sequence reads were trimmed and mapped using CLC Genomics Workbench (CLC Bio, Denmark) using standard parameters. Subsequently, inversion, deletions, structural variants, and single nucleotide polymorphisms were determined using standard parameters.

### Subcommunity enrichment

Forty-eight growth conditions were established (Supplementary Table X). Drug compounds, except rifaximin which was freshly prepared, were dissolved as 100x stock solutions in ethanol, DMSO, or water, as appropriate, and stored at −20 °C in 96-well drug plates. Rich and selective media were freshly prepared and stored at room temperature. Twenty-four hours prior to inoculation, all materials and reagents were transferred into an anaerobic chamber (Coy Laboratories). On the day of inoculation, drug/medium plates were assembled under anaerobic conditions by mixing 100 µl of medium with the compounds in the drug plate, followed by transfer of 115 µl of the resulting drug–medium mixture back into the deep-well plate. For medium-no drug conditions, 1,485 µl of medium was added to the corresponding well.

Cryopreserved stool from donor MB003 (700 µl aliquot, prepared as described above) was diluted 1:100 in PBS, and 15 µl of this suspension was inoculated into each of the 48 wells of a 96-well deep-well plate containing 1,485 µl of medium (∼5 × 10⁷ CFU ml⁻¹ final). Cultures were incubated anaerobically at 37 °C for 24 h, then sequentially passaged for three additional days by 1:50 dilution into fresh drug/medium plates. Growth was monitored daily by transferring 100 µl of culture into shallow 96-well plates and measuring optical density at 578 nm every hour for 24 h. On day 5, 500 µl of each culture was mixed with glycerol for long-term storage, and the remaining culture (∼1 ml) was collected for DNA extraction.

*E. coli* was isolated from the bile-salt-treated subcommunity by plating on LB agar plates and incubating at 37°C under aerobic conditions overnight. Colonies were purified and species was determined by 16S sequencing. After confirmation, that the isolated *E. coli* strain is selective for *K. pneumoniae ompK^evo^*, whole genome sequencing of the strain was performed as described previously.

### Construction of knockout mutants

Gene knockouts were constructed using the lambda red recombineering system encoded on the synthetic vector pSIM5-tet ^60,61^. Overnight cultures of the strain of interest were diluted 1:100 fold in 50 ml LB broth at 37°C with constant shaking to a final OD600 of 0.2. Cells were harvested and washed three times with ice-cold 10% glycerol and taken up in a final volume of 100µl. 40µl cells were mixed with 50 ng of pSIM5 tet and transferred to an electroporation cuvette (1-mm gap). Electroporation was performed using a Gene Pulser (Bio-Rad) at 1.9kV, 400 Ohm and 25µF. After electroporation, cells were recovered in 1ml SOC (2% trypton, 0.5% yeast extract, 10 mM NaCl, 2.5 mM KCl, 10 mM MgCl_2_, 10 mM MgSO_4_ and 20 mM Glucose) for 2 hours, plated on LB Agar plates containing 10 mg/L tetracycline and incubated at 30°C.

To prepare electrocompetent cells, pSIM5-tet carrying strains were grown overnight in LB broth supplemented with 10 mg/L tetracycline at 30°C, diluted 1:100 in 50 ml LB broth supplemented with 10 mg/L tetracycline and incubated at 30° until they reached OD600 = 0.02. Cultures were moved to a 42°C water-bath to induce expression of the lambda red system for 15 minutes. Afterwards, the cultures were placed on ice for 10 minutes and washed 3 times with ice-cold 10% glycerol. The cells were resuspended in 200µl 10% glycerol final volume, and 40µl were mixed with a 200ng purified PCR product of a gene construct containing *amilCP* (blue chromoprotein), *cat* (chloramphenicol resistance) and *sacB* (sucrose sensitivity) with homologous overhangs up- and downstream of the gene of interest. Electroporation and cell recovery were performed as described above. Successful transformants were selected on LB agar plates supplemented with 15 mg/L chloramphenicol at 30°C. Transformants were restreaked on LB agar plates supplemented with 10 mg/L tetracycline at 30°C to ensure maintenance of pSIM5-tet. The strains were then subjected to a second round of transformation as described above. Here, instead of a PCR product, single-stranded DNA was electroporated composed only of the up- and down-stream homologous overhangs, to replace the amilCP-cat-sacB cassette leaving a scarless knockout. Recovery of transformants was performed in 5 ml SOC overnight to ensure dilution of the SacB protein in cells, and successful transformants were selected on salt-free LB agar plates supplemented with 5% sucrose. Loss AmilCP, and therewith loss of the blue color, was an additional verification, that the complete amilCP-cat-sacB cassette was removed, and sucrose sensitivity was not only lost due to a loss-of-function mutation in *sacB*. All final constructs were confirmed by Sanger sequencing.

### Growth characterization in spent medium

To prepare spent medium an overnight culture of the E. coli strain of interest was diluted 1:1000 in 20 ml mGAM under anaerobic conditions. The culture was grown for 20h, and cells were pelleted by centrifugation for 10 minutes at 4000 rpm. The supernatant was transferred and subjected to a second round of centrifugation. The supernatant of this second round was sterile filtered using a filter with a 0.22µm pore size. The filtered supernatant was brought back to anaerobic conditions and used as ‘spent medium broth’ for growth characterization of the *K. pneumoniae* isolates. For this, overnight cultures of K. pneumoniae in mGAM were diluted 1:2000 in spent medium. Growth rate was determined in 96-well plates sealed with breathable membranes and OD600 was determined in 20-minute intervals over 24hours using a Filtermax F5 multimode plate reader (Molecular Devices). The first timepoint was used for background subtraction. All growth rate determinations were performed in triplicate.

### Estimating the occurrence and frequency of SNPs across the *glyP* operon among *K. pneumoniae* CC258 genomes

We generated a large collection of CC258 genomes from publicly-available genomes available in Pathogenwatch (https://pathogen.watch)^62^. Raw sequence reads associated with these genomes were downloaded from the European Nucleotide Archive (ENA) using available accession numbers. Reads were mapped to the ST11 reference genome, HS11286 (accession CP003200) using a workflow (https://gitlab.com/cgps/ghru/pipelines/snp_phylogeny) (version 1.2.2) incorporating BWA-MEM (Li, 2013), SAMtools and BCFtools^63^ and a pseudo-genome alignment was generated. Genomes were excluded from downstream analyses if they had < 20x average mapping depth or ≥ 25% missing sites in the pseudo-genome alignment. Gubbins v3.2.1^64^ was used to remove recombined regions from the alignment and generate a maximum likelihood tree with RAxML-NG^65^. For each base position within the pseudo-genome alignment, a maximum parsimony method was used to perform ancestral reconstruction across the tree (https://github.com/sanger-pathogens/bact-gen-scripts/blob/master/reconstruct_snps_on_tree.py). The number of state changes inferred to have occurred across the tree at each base position was calculated and the types of SNPs recorded.

### Genotypic characterization of genome assemblies

Genome assemblies from the above-described CC258 collection were additionally characterized. Kleborate v2.3.0 was used to identify virulence and resistance genes including alterations within the *ompK35* and *ompK36* porin genes^66^. These included truncations of both genes, and loop 3 insertions (glycine-aspartate (GD) or threonine-aspartate (TD)) within *ompK36*. The *lacI* type transcription factor was identified among the assemblies using BLASTn v2.14.1^67^ with the full-length gene from ATCC43816 (accession CP009208) as the query sequence. We required a single hit in each assembly that matched ≥25% of the query length and contained a start codon to unambiguously identify the gene. Nucleotide sequences were translated to protein sequences using Seaview v5.0.5^68^. Protein sequences were predicted to be intact if they possessed at least 90% of the query length. Metadata and genotypic data were visualised with the phylogenetic trees using Microreact v273^69^.

The Artemis Comparison Tool (ACT) v18.1.0^70^ was used to compare the ST11 genome, HS11286, with a representative genome assembly from the clade of 306 CC258 isolates with an absent lacI type TF gene (SRR12509433). Prior to the comparison, the contigs from SRR12509433 were aligned and reordered against the HS11286 complete genome using Abacas v1.3.1^71^.

### Metabolite extraction

LC-MS grade solvents were purchased from Fisher Scientific, and all other chemicals from Sigma Aldrich, unless otherwise specified. For metabolite extraction, 20 µL of the bacterial supernatants were transferred in a 96-well plate with 105 µL of ACN:MeOH solvent mixture in a 1:1 ratio supplemented with an internal standard mixture. The internal standard mixture contained sulfamethoxazole, caffeine, ipriflavone, and warfarin, each at 80 nM final concentration. Following, the samples were chilled at -20°C for 1 h to allow proteins to precipitate, then centrifuged at 4500 rpm for 15 min at 4 °C. The remaining supernatant was finally aliquoted in 96-well microtiter plates, diluted with water (1:1), and an additional centrifugation step at 4500 rpm for 15 mins and 4 °C.

### LCMS analysis

LC-MS analysis was performed using reverse-phase chromatography in negative ionization mode. Chromatographic separation was performed using an InfinityLab Poroshell 120 HILIC-Z column (2.1 x 150 mm, 2.7-micron pore size) in an Agilent 1200 Infinity UHPLC system and mobile phases A: H_2_O + 5mM Ammonium formate + 0.1% formic acid and B: acetonitrile + 5mM Ammonium formate + 0.1% formic acid. The column compartment was kept at 25°C. 5 µL of the sample was injected at 98% B and 0.250 mL/min flow followed by: 98% B until minute 3, a gradient to 70% B until minute 11, a gradient to 60% B until minute 12, a gradient to 5% B to minute 16, remaining at 5% until minute 18 before re-equilibration to 98% until minute 20. Flow remained at 0.250 mL/min.

The qTOF (Agilent 6550) was operated in positive scanning mode (50 – 1500 *m/z*) and electrospray ionization (ESI) with the following source parameters: VCap: 3000 V, nozzle voltage: 0 V, gas temp: 225 °C; drying gas 11 L/min; nebulizer: 40 psig; sheath gas temp 225°C; sheath gas flow 10 L/min; fragmentor 300V and Octopole RF Vpp 450V. Online mass calibration was performed using a second ionization source and a constant flow of reference solution (121.0508 and 922.0097 *m/z*). Tandem mass spectrometry analysis (LC-MS/MS) was performed for all significant metabolites using the chromatographic separation and source parameters described above and the targeted-MS/MS mode of the instrument with a preferred inclusion list for parent ion with 20 ppm tolerance, Iso width set to ‘narrow width’, and collision energy set to either 10, 20, or 40 eV.

### Data Extraction and Statistical Analysis

To reduce data complexity for downstream analysis, mass spectrometry data were centroided and converted from the proprietary format (.d) to the m/z extensible markup language format (.cef) using the peak picking algorithm of Agilent DAReprocessor at the following parameters: minimum peaks height >= 5.000 counts; all ion species; only select common organic compounds (no halogens); isotoping grouping with limit assigned charge states to a range of 1; compound filters based on Quality Score > 70. The .cef converted data represents each peak in the mass-to-charge (*m/z*) dimension as a single, discrete peak in the mass spectrum, reducing data size and the grouping of adducts for every molecular feature.

Following, the table of putative metabolic features, along with their relative intensities, was then aligned with Agilent Mass Profiler Professional - MPP (version 15.1). In brief, feature alignment allowed mass tolerances of 0.003 AMU or 20 ppm and retention time tolerance of 0.3 min or 3%; minimum absolute abundance 10.000 counts; minimum number of ions 3; multiple charge states forbidden. The final feature table was also filtered to retain only features that appear in 75% of replicates of at least one condition and submitted to MS1 feature annotation against the Metabolite and Tandem MS Database (METLIN) (v1233)^72^ was performed by Agilent MPP. Human Metabolome Database (HMDB) accession numbers were also derived from METLIN, and compound classes were assigned based on the latest HMDB release (v5.0)^73^.

The resulting feature quantification table was processed in R (v4.4.1) using RStudio/2024.09.1, applying an adapted protocol for blank removal, imputation, quantile normalization, and log- transformation^74^. In brief, features with specific blank-to-sample ratios and samples with abnormally low mean intensities were flagged as poor-quality and removed from analysis. The remaining data were imputed to address missing values, in which a limit of detection (LOD) threshold was established based on the minimum non-zero intensity in the data. Values below this threshold were imputed to the LOD cutoff. Intensities were then quantile-normalized using the normalize.quantiles function from the preprocessCore package (v1.64.0)^75^ and subsequently log₂ transformed with a pseudo-count (half of the minimum positive value) to stabilize variance was applied. For multivariate analysis such as PCA, data were also z-scored (mean-centering and unit-variance scaling).

Multivariate and statistical analyses were performed in R 4.4.1 using RStudio/2023.06.1. First, the prcomp function was used for Principal Component Analysis (PCA) for data visualization, detection of sample outliers, and overall sample similarity. Volcano plots were done to statistically compare spent media before and after Kpn inoculation. The significance of a molecular feature was based on the intensity differences, in which the p-values were assessed by t-test (t.test function in R). The p-values were FDR-corrected for multiple hypotheses testing using the Benjamini-Hochberg procedure (p.adjust function in R with ‘fdr’ parameter). Significant metabolites were selected when their intensity had a fold change > 0.5 and a corrected p-value < 0.05. Finally, the area under the curve from significant features was integrated using the MassHunter Quantitative Analysis Software (Agilent, version 7.0) based on the accurate high-resolution mass and RT of the reference analytes with the following parameters: signal threshold of 30,000; mass tolerances of 0.003 amu or 20 ppm; retention time tolerance of 0.5 min. All metabolites annotated in this manuscript were assigned a Level 1 annotation according to the Metabolomics Standards Initiative (MSI) classification system. This includes a match in retention time using the same chromatographic system, accurate mass, and MS/MS fragmentation pattern, ensuring unequivocal identification of the compounds.

### Proteomics

For the differential abundance proteomics, cultures were grown anaerobically at °C to mid log phase (OD600 = 0.5), washed once with PBS and snap frozen. Pellets were resuspended in 2% SDS and incubated at 99 °C for 5 min.

TPP was performed as previously described^46,76^. In brief, *K. pneumoniae ompK^evo^* cultures were grown anaerobically at 37 °C overnight. Cells were then washed twice with anaerobic PBS, resuspended in lysis buffer (50 ug/ml lysozyme, 1mM MgCl2, cOmplete protease inhibitors, 0.25 U/ul benzonase in PBS) and lysed with 5 freeze-thaw cycles. Crude lysates (protein concentrations normalized to 10 ug/ul) were treated with filter-sterilized spent medium prepared from various *E. coli* strains as described previously for 10 min at RT. Aliquots of treated lysates were then incubated in 10 different temperatures for 3 min, followed by 3 min at RT and 5min on ice. NP-40 was added to a final concentration of 0.8%, aggregates were removed by filtration and soluble proteins were prepared for mass spec.

Proteins were prepared and digested as previously described^47,77^. Eluted peptides were labelled with TMT16plex, for the differential abundance proteomics, and TMT18plex, for the TPP experiment. Samples were then pooled (all samples of each two adjacent temperatures for the TPP experiment) and pre-fractionated into 6 fractions under high pH conditions. Samples were then analyzed with liquid chromatography coupled to tandem mass spectrometry, as previously described^46^.

For the differential abundance proteomics, MS data were processed as previously described^47^. Briefly, raw MS files were processed with isobarQuant^78^ and the identification of peptide and protein was performed with Mascot 1.0 (Matrix Science) against a FASTA database of translated ORFs annotated in CP009208 modified to include known contaminants and the reversed protein sequences. For the TPP experiment, raw files were converted to mzML format using MSConvert from ProteoWizard, using peak picking, 64-bit encoding and zlib compression, and filtering for the 1000 most intense peaks. Files were then searched using MSFragger (v4.0)^79^ in FragPipe (21.1) against a FASTA database of translated ORFs annotated in CP009208 containing common contaminants and reversed sequences.

Data analysis was performed using R. Proteins with at least one uniquely identified peptide were kept for the analysis. For the differential abundance proteomics, protein intensities across all strains were normalized using variance stabilization normalization^80^ and limma was used to assess the differential expression of proteins. For the TPP experiment, protein intensities across all conditions within each temperature were normalized as above and the abundance and thermal stability scores were calculated as previously described ^47,81^. To assess the significance of abundance and thermal stability scores, limma was used ^82^.

## Supplementary material

Extended Data Table 1 – Whole proteome abundance changes in the *K. pneumoniae ompK^evo^* clones versus the wild type

Extended Data Table 2 – Whole proteome thermal stability changes Whole proteome abundance changes in the *K. pneumoniae ompK^evo^* clones versus the wild type in presence of *E. coli^MB003^* spent medium

## Data availability

Sequencing data of the *K. pneumoniae* isolates has been deposited in the European Nucleotide Archive (ENA) at EMBL-EBI under accession number PRJEB110171. *K. pneumoniae* CC258 genomes used in the ancestral reconstruction analysis are publicly-available with accessions included in the Microreact project: https://microreact.org/project/kp-cc258. The mass spectrometry proteomics data have been deposited to the ProteomeXchange Consortium via the PRIDE partner repository with the dataset identifier PXD075097. The metabolomics data is available in the MetaboLights database (https://www.ebi.ac.uk/metabolights/) under accession number MTBLS7388.

## Acknowledgements

We thank A. Mateus for assistance with proteomic analyses, D. Hazenbrink and C. G. P. Voogdt for assistance with subcommunity enrichments of fecal microbiomes, Maria I. Keller for DNA extraction of fecal microbiome samples and S. Giri for valuable discussions throughout the project. We also thank the EMBL core facilities for Genomics, Metabolomics, Proteomics, Flow Cytometry and Microbial Automation & Culturomics for their extended support throughout the project.

## Author contribution

M.K. and A.T. acquired funding and conceptualized the study. M.K., S.G-S., L.M, D.P., V.C. and D.M.S. performed the experiments. S.D., J.L.C.W. and N.K. performed the computational analysis. G.F., M.Z., M.M. and A.T. supervised this study. M.K., L.M., D.P., S.D. and N.K. visualized the data and generated the figures. M.K. and A.T. wrote the original draft of the manuscript. All authors discussed the results and reviewed and edited the final version of the manuscript.

---

**Extended Data Fig. 1:**
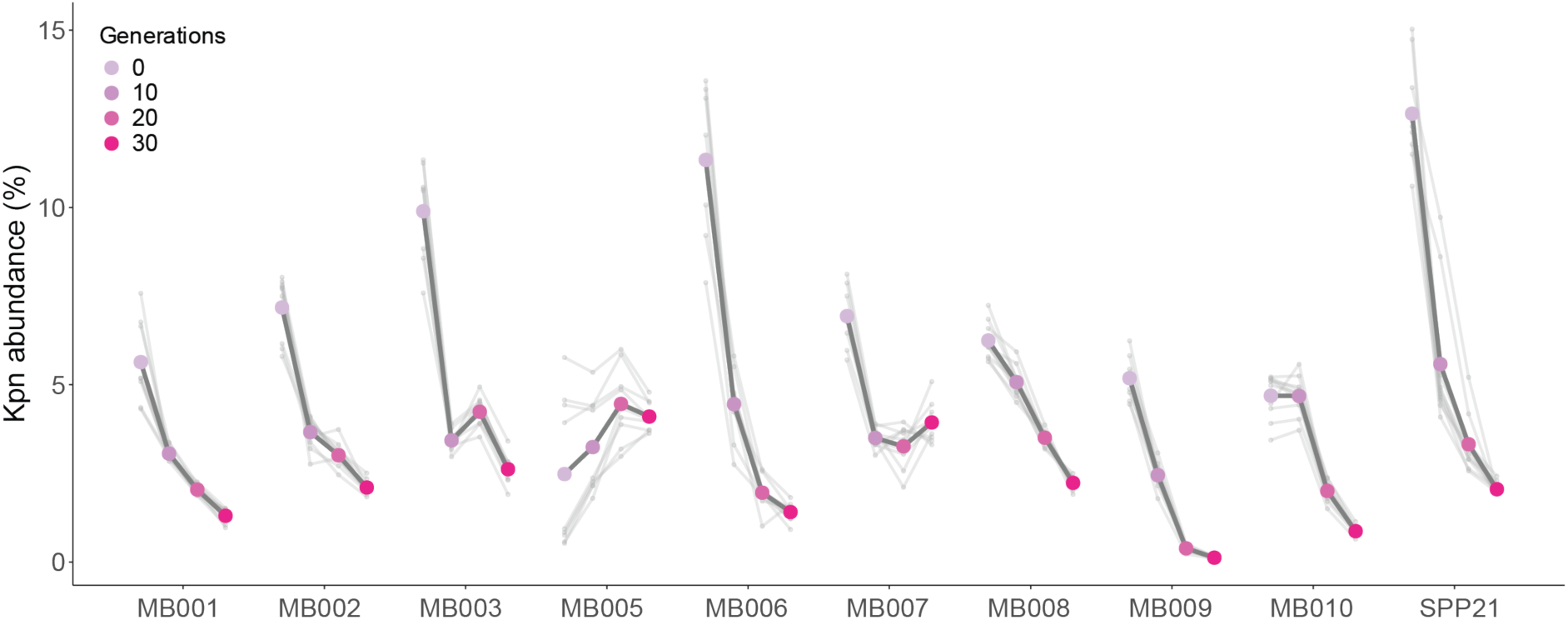
Abundance of *K. pneumoniae* wildtype in different microbiomes over the course of the competition experiment. Dark grey lines show the mean total abundance of the *K. pneumoniae* competitors in various fecal derived and synthetic microbiomes, whereas light grey lines indicate abundance in each replicate. Abundancies were determined by flow cytometry, where GFP^+^ and RFP^+^ cells represented *K. pneumoniae* and nonfluorescent events represent community members. Competitions were passaged for 4 days in mGAM under anaerobic conditions at 37 °C with a daily dilution factor of 1:1000 corresponding to about 10 generations of growth per day.

**Extended Data Fig. 2:**
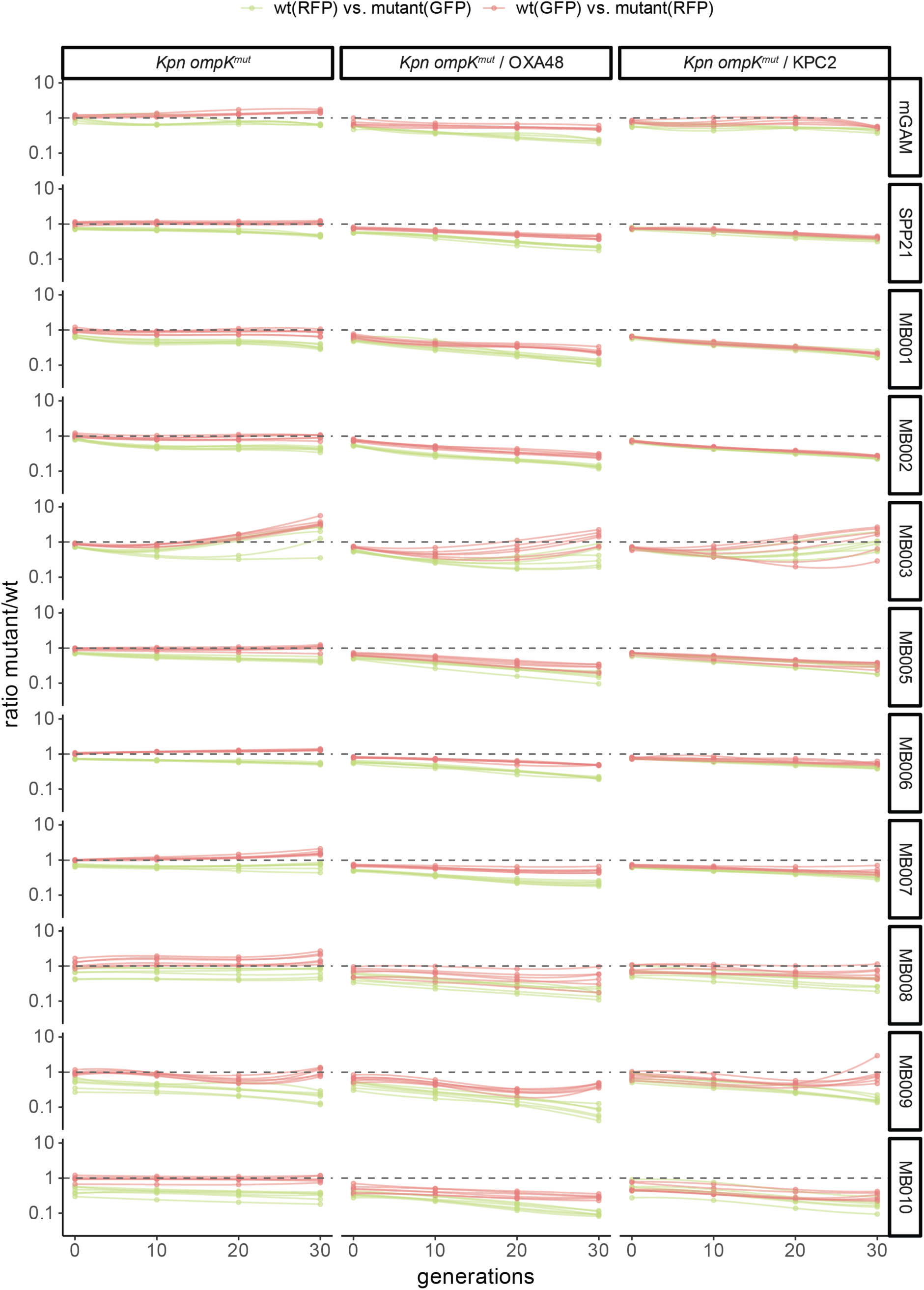
Change in ratios over time of *K. pneumoniae* competitors in presence and absence of various communities. Green and red data points represent competitions where the resistant mutant (*K. pneumoniae ompK^mut^, ompK^mut^* /pOXA48 *and ompK^mut^* /KPC2) was tagged with GFP or RFP, respectively. Ratios were determined daily by flow cytometry. Competitions were performed as described in Extended Data Fig. 1.

**Extended Data Fig. 3:**
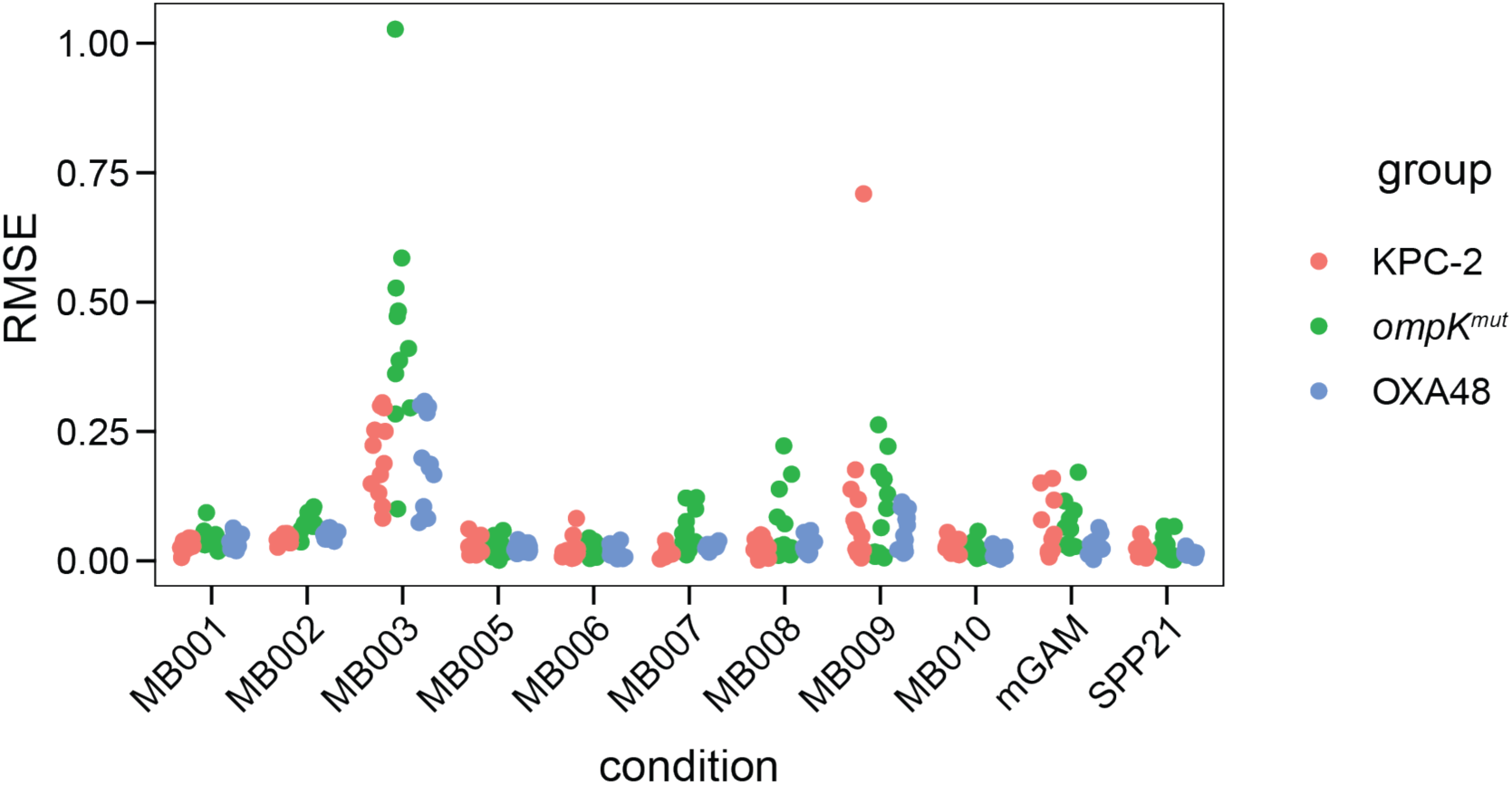
Fit measures for selection coefficients. Root mean squared error (RSME) of linear models fits of change in ratios of indicated *K. pneumoniae* strain over the wildtype as a function of time in different community compositions. Each datapoint represents the RSME of an individual replicate of a longitudinal fitness experiment. Corresponds to curves shown in Extended Data Fig. 2

**Extended Data Fig. 4:**
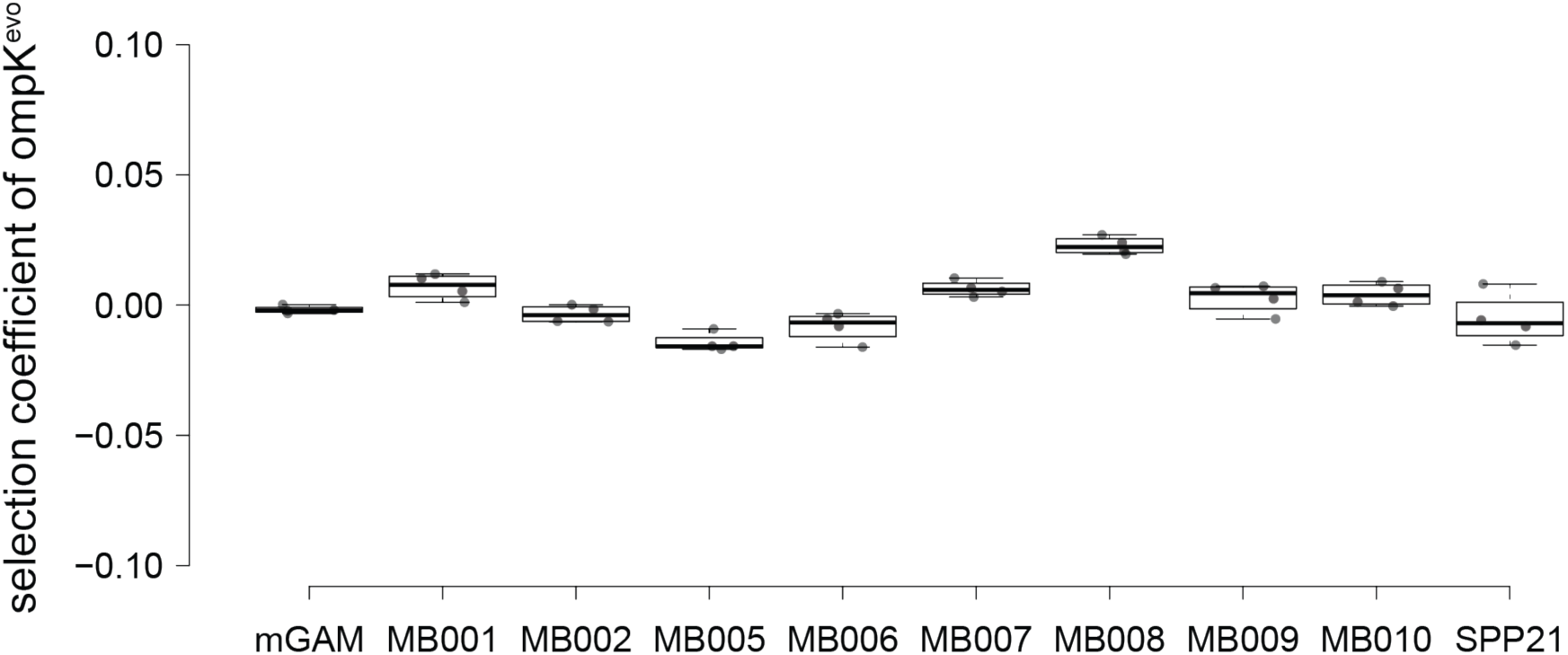
Selection coefficient of *K. pneumoniae ompK^evo^* in different microbiomes. *K. pneumoniae ompK^evo^* GFP^+^ was competed with an RFP^+^ *K. pneumoniae* wildtype over 4 days using the previously described competition setup in presence or absence of different microbial communities. Box plots are depicted as in Fig. 1g (n is minimum of four biological replicates).

**Extended Data Fig. 5:**
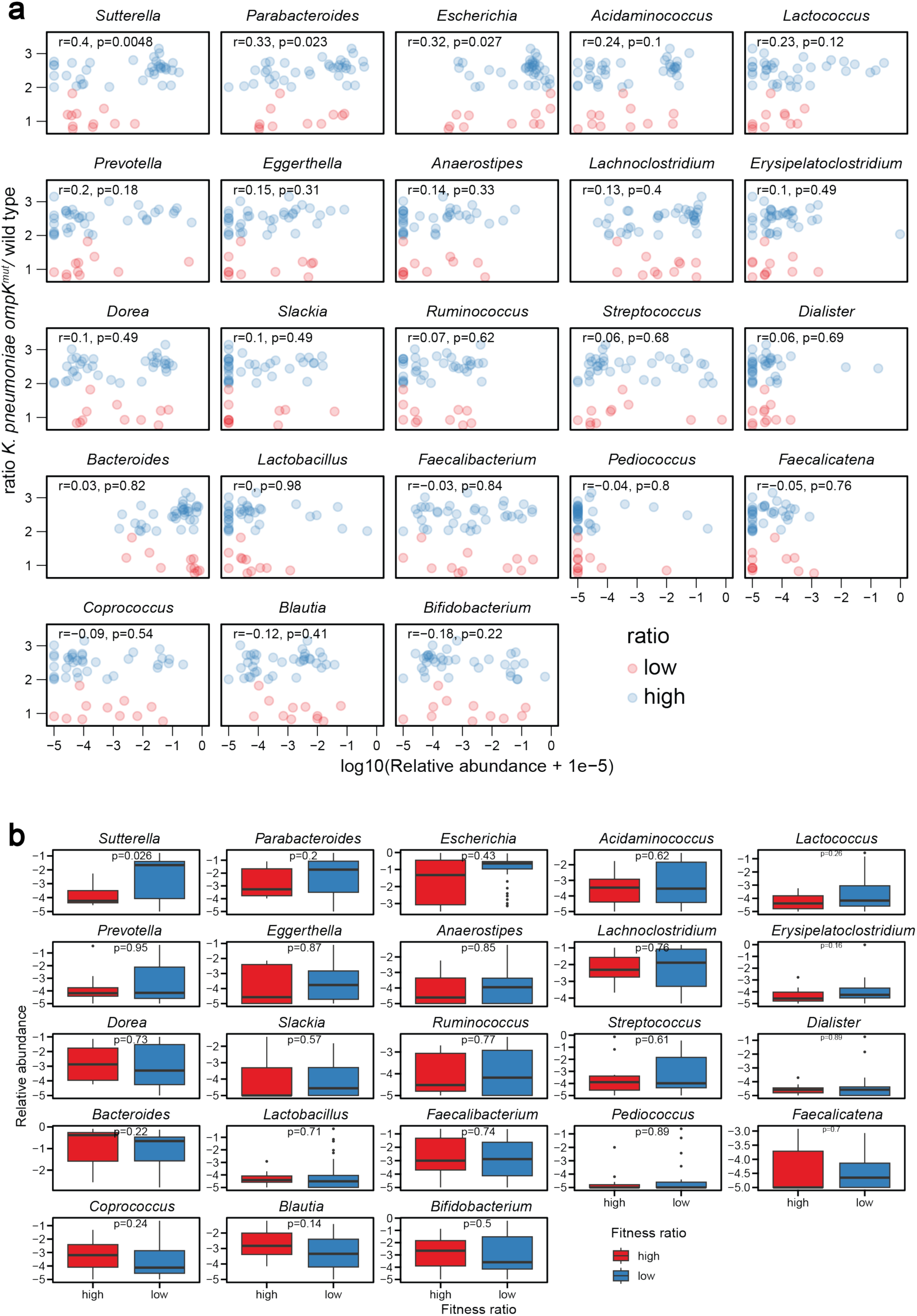
MB003 subcommunity selection of *K. pneumoniae ompK^evo^* over *K. pneumoniae*. a, Scatter plot of *K. pneumoniae ompK^evo^* ove*r* wildtype ratios against taxon relative abundances in microbiome MB003 derived subcommunities. Relative abundances of taxa are log10-scaled. Conditions are grouped into high and low fitness ratios, where conditions that enriched *K. pneumoniae ompK^evo^* ove*r* wildtype at least 2-fold are considered high, everything below as low. P-values were computed using a Pearson correlation test. Only genera with a relative abundance of at least 5% in at least one of the perturbation conditions are shown. **b,** Box plot of taxon relative abundances between high fitness ratio and low fitness ratio perturbation conditions. Relative abundances of taxa are log10-scaled. Conditions are grouped into high and low fitness. P-values were computed using a Wilcoxon rank-sum test. Only genera with a relative abundance of at least 5% in at least one of the perturbation conditions are shown.

**Extended Data Fig. 6:**
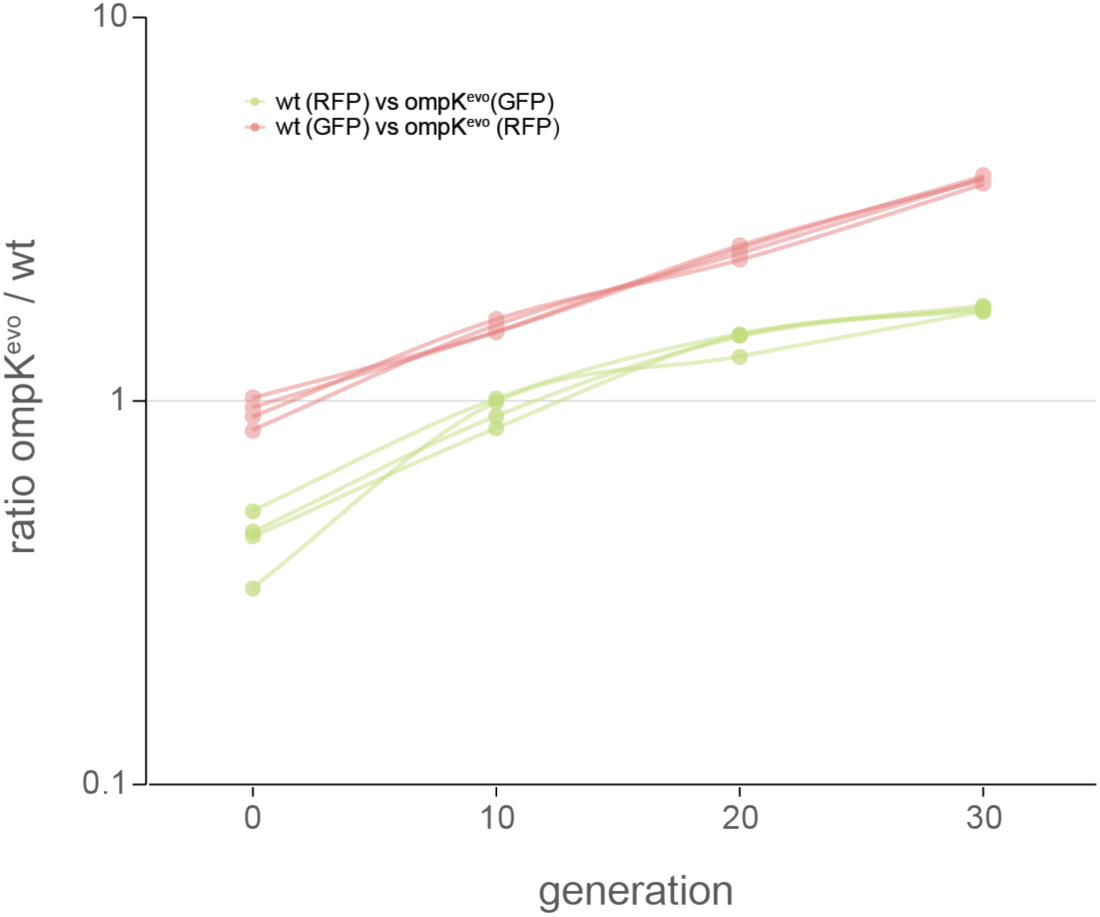
Ratio of *K. pneumoniae ompK^evo^* over the wildtype in the presence of *E. coli^MB003^* over 40 generations of growth. Each line represents a replicate competition and colors indicate the marker carried by the porin mutant (green = GFP, red = RFP)

**Extended Data Fig. 7:**
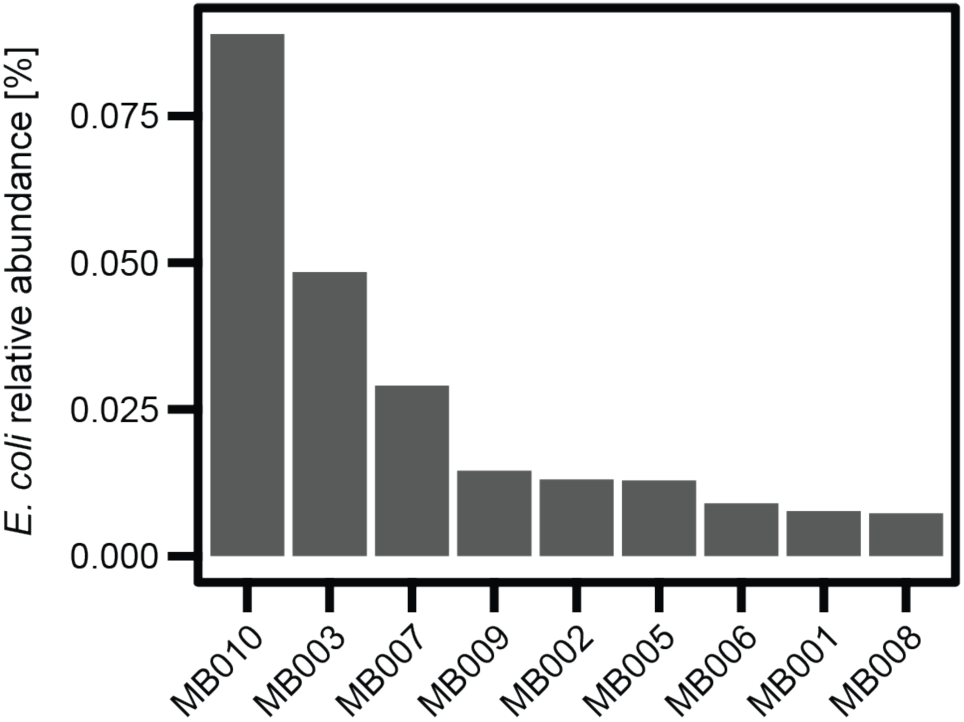
Abundance of *E. coli* in native stool samples from MB001-MB010. This is based on metagenomics analysis of MB001-MB010 shown in Fig. 2a.

**Extended Data Fig. 8:**
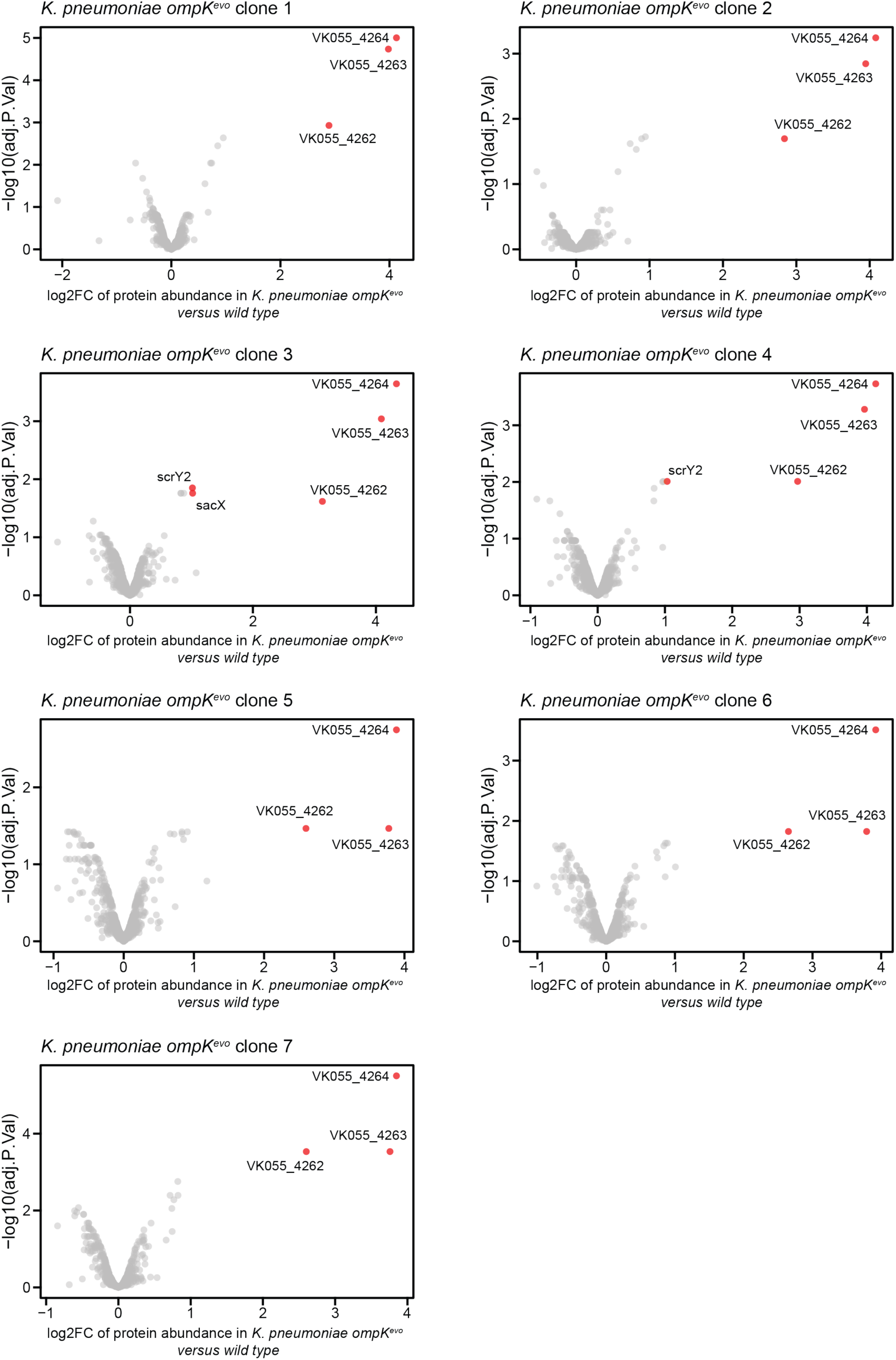
Protein abundance changes in *K. pneumoniae ompK^evo^* versus wildtype. The volcano plots show the up- or downregulated proteins in all seven isolated *K. pneumoniae ompK^evo^* clones. Cells were grown overnight in mGAM under anaerobic conditions and harvested after 20h for whole cell proteome analysis using mass spectrometry. Each strain was tested in three biological replicates.

**Extended Data Fig. 9:**
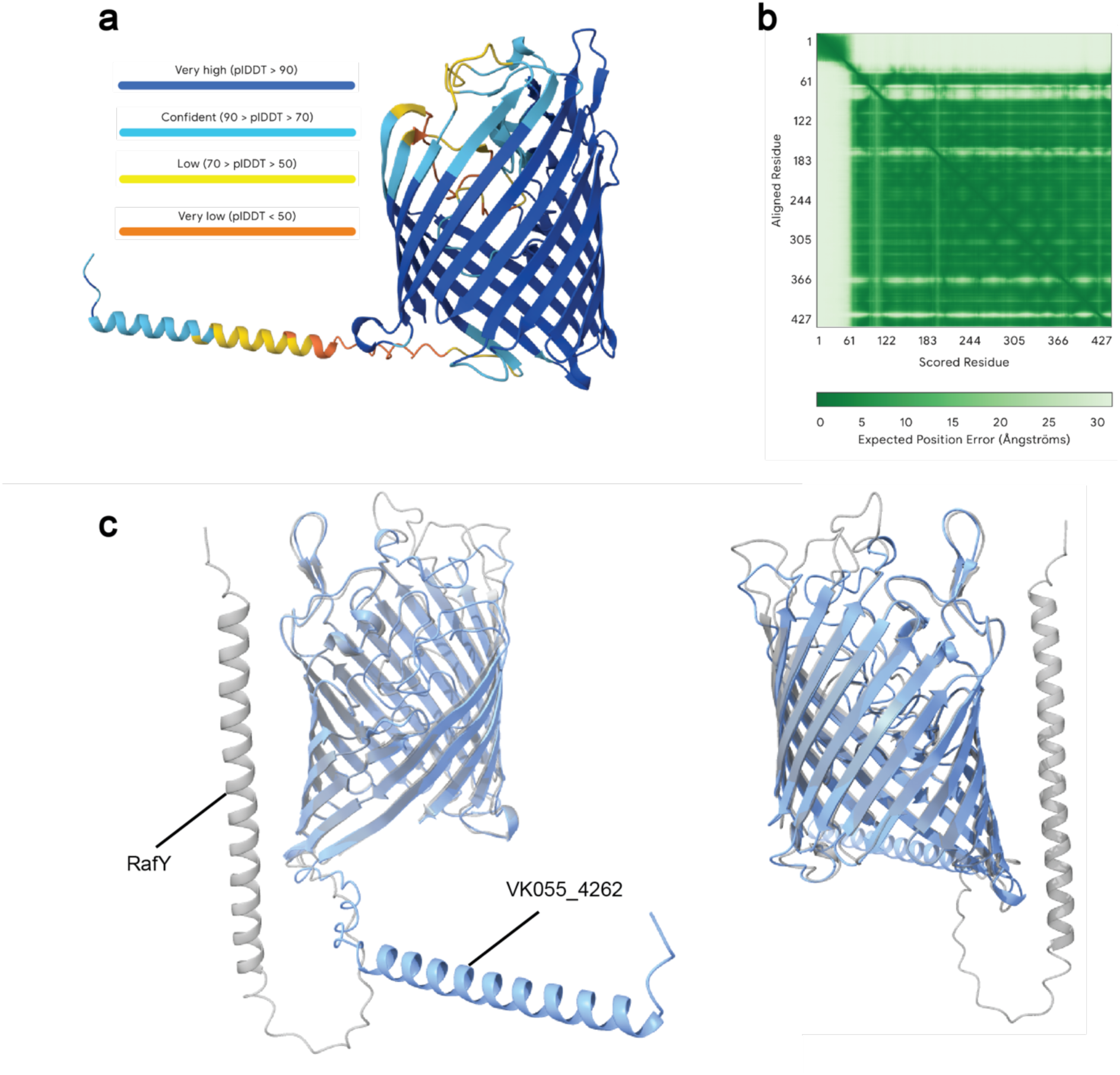
Structural prediction and comparisons for GlyP. **a**, AlphaFold prediction of the *K. pneumoniae* porin upregulated in *K. pneumoniae ompK^evo^*. The AlphaFold-predicted structure of the mature protein (signal peptide removed) is shown as a cartoon representation colored by pLDDT confidence scores. The ß-barrel core is predicted with high confidence. **b**, Predicted aligned error (PAE) plot for the AlphaFold model. The PAE plot indicates high confidence in the relative positioning of residues withing the ß-barrel core, while increased PAE is primarily associated with extracellular loop regions. The short cytoplasmic helix also displays reduced confidence, consistent with local flexibility and disorder. **c,** Structure alignment of GlyP with RafY. UCSF ChimeraX structural superposition of the AlphaFold-predicted mature porin (blue) and RafY (grey) shows close alignment of the ß-barrel core with divergence in loop regions. The alignment produced a sequence alignment score of 1495.9 and an RMSD of 0.67Å over 312 pruned atom pairs (22.3Å across all aligned residues).

**Extended Data Fig. 10:**
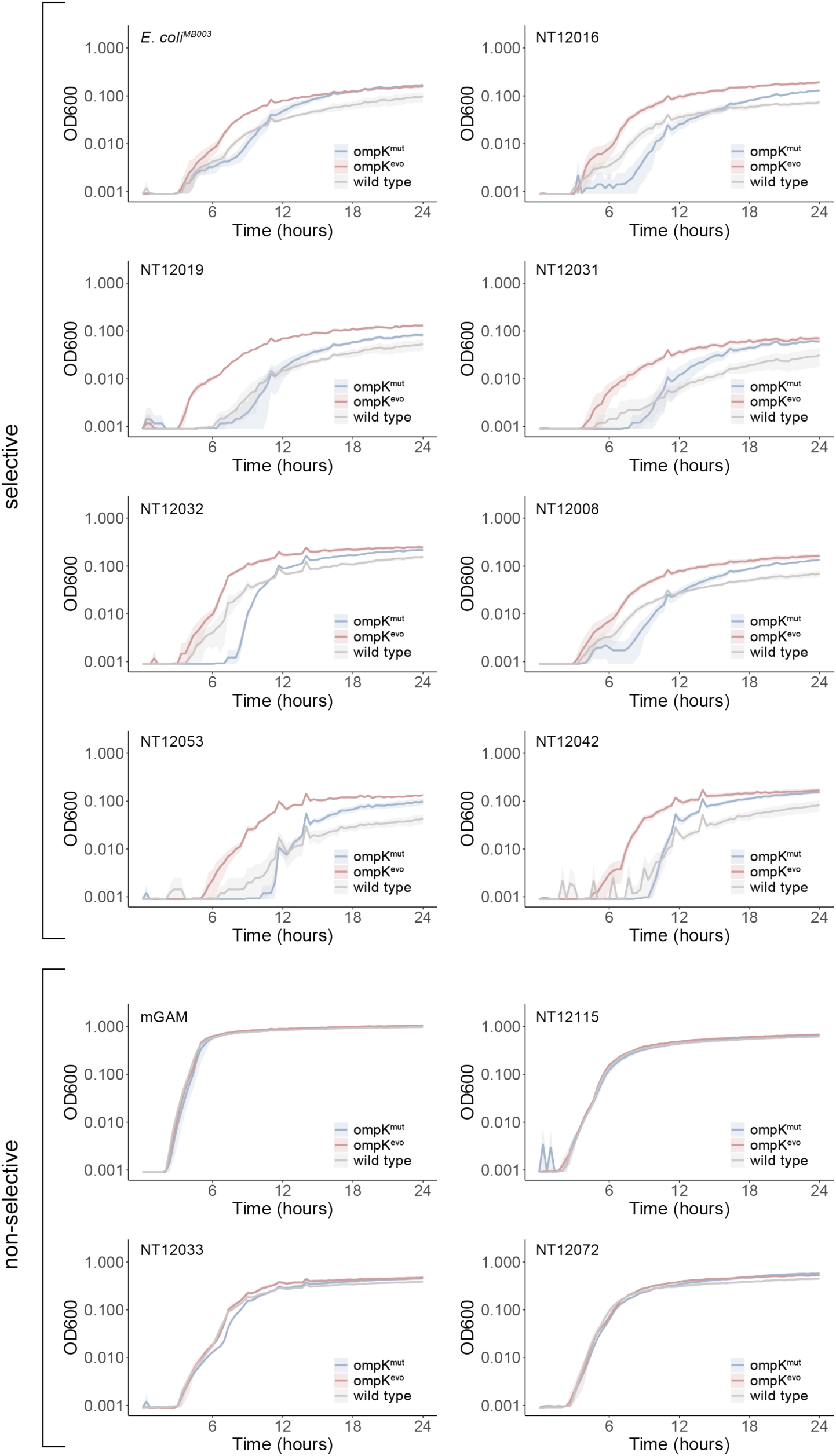
Growth rates of *K. pneumoniae wildtype, ompK^mut^ and ompK^evo^* in various *E. coli*-spent media. The spent media were prepared by growing different *E. coli* strains overnight in mGAM at 37 °C under anaerobic conditions and subsequent sterile filtration using a 0.22µM pore size filter. Lines represent average of four independent replicates, shades represent the standard deviation.

**Extended Data Fig. 11:**
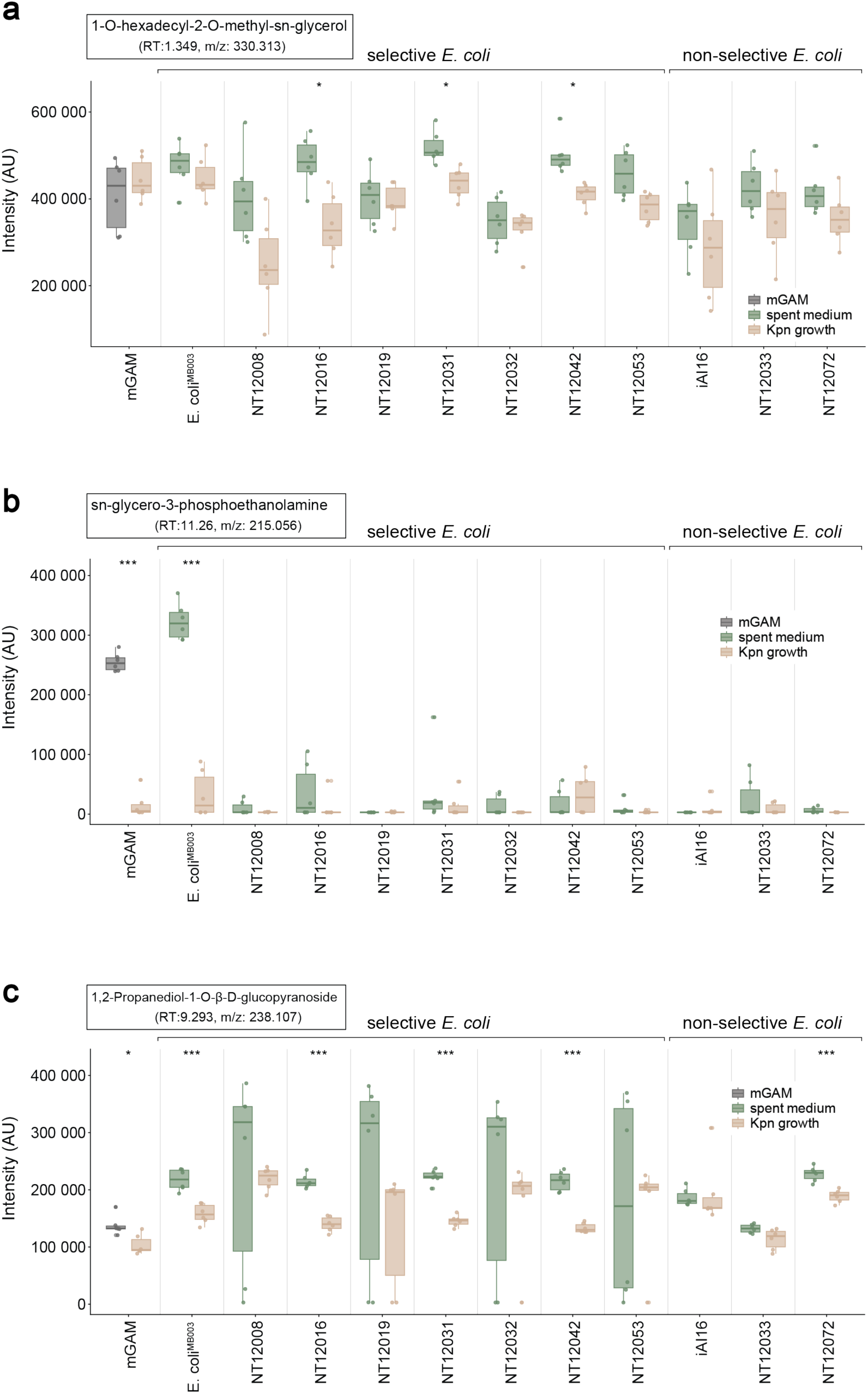
Abundance of glycerol-containing metabolites in *E. coli*-spent media in the presence or absence of *K. pneumoniae*. Absolute LC-MS signal intensities of (a) 1-O-hexadecyl-2-O-methyl-sn-glycerol (m/z 330.313), (b) sn-glycero-3-phosphoethanolamine (m/z 215.059) and (c) 1,2-Propanediol-1-O-β-D-glucopyranoside (m/z 238.107) in culture supernatants of *E. coli* before and after growth of *K. pneumoniae ompK^evo^*. A reduction in signal intensity indicates depletion of these metabolites. Values are based on 6 biological replicates. Statistical significance was assessed using a t-test. * p_adjusted_ < 0.05, ** p_adjusted_ < 0.01, *** p_adjusted_ < 0.001

